# Developmentally dynamic changes in DNA methylation in the human pancreas

**DOI:** 10.1101/2023.10.19.563104

**Authors:** Ailsa MacCalman, Elisa De Franco, Alice Franklin, Christine S. Flaxman, Sarah J. Richardson, Kathryn Murrall, Joe Burrage, Barts Pancreas Tissue Bank (BPTB), Emma M Walker, Noel G. Morgan, Andrew T. Hattersley, Emma L. Dempster, Eilis J. Hannon, Aaron R. Jeffries, Nick D. L. Owens, Jonathan Mill

## Abstract

Development of the human pancreas requires the precise temporal control of gene expression via epigenetic mechanisms and the binding of key transcription factors. We quantified genome-wide patterns of DNA methylation in human fetal pancreatic samples from donors aged 6 to 21 post-conception weeks. We found dramatic changes in DNA methylation across pancreas development, with >21% of sites characterized as developmental differentially methylated positions (dDMPs) including many annotated to genes associated with monogenic diabetes. An analysis of DNA methylation in postnatal pancreas tissue showed that the dramatic temporal changes in DNA methylation occurring in the developing pancreas are largely limited to the prenatal period. Significant differences in DNA methylation were observed between males and females at a number of autosomal sites, with a small proportion of sites showing sex-specific DNA methylation trajectories across pancreas development. Pancreas dDMPs were not distributed equally across the genome, and were depleted in regulatory domains characterized by open chromatin and the binding of known pancreatic development transcription factors. Finally, we compared our pancreas dDMPs to previous findings from the human brain, identifying evidence for tissue-specific developmental changes in DNA methylation. To our knowledge, this represents the most extensive exploration of DNA methylation patterns during human fetal pancreas development, confirming the prenatal period as a time of major epigenomic plasticity.

## INTRODUCTION

Development of the human pancreas is a highly intricate process requiring the precise temporal control of gene expression. The mature pancreas derives from two buds (dorsal and ventral) originating from the distal foregut endoderm and has both exocrine and endocrine functions reflecting distinct cellular lineages (1). The exocrine pancreas, responsible for the secretion of digestive enzymes, is composed principally of acinar and ductal cells. The endocrine pancreas, assembled into Islets of Langerhans, contains α-cells, β-cells, δ-cells, ε-cells, and γ-cells (otherwise known as PP-cells) which each release specific hormones (glucagon, insulin, somatostatin, ghrelin and pancreatic polypeptide, respectively) that act to regulate blood glucose levels (2). The diversity of the cells within the maturing pancreas indicates a complex developmental pathway involving spatially-and temporally-coordinated changes in gene regulation (3), although the precise molecular mechanisms driving this process are not fully understood. Importantly, several diseases are known to result from aberrant development of the pancreas including maturity onset diabetes of the young (MODY) and neonatal diabetes - monogenic disorders caused by pathogenic mutations in genes involved in coordinating pancreas development (4, 5).

Epigenetic modifications play a key role in the dynamic regulation of gene function during tissue development and differentiation (6). The most extensively studied epigenetic mechanism is DNA methylation, the covalent addition of a methyl group to the fifth carbon position of cytosine (7). DNA methylation is traditionally considered to repress local gene expression via disruption of transcription factor binding and the recruitment of methyl-binding proteins that initiate chromatin compaction and gene silencing (8). Evidence suggests there is a more nuanced relationship between DNA methylation and transcription that is dependent on genomic and cellular context; DNA methylation in the gene body, for example, is often associated with increased expression (9) and has been implicated in other genomic functions including alternative splicing and promoter usage (10).

The establishment and maintenance of tissue-specific DNA methylation patterns is a crucial feature for mammalian development, and within the pancreas, DNA methylation plays a critical role in the specification of the endocrine and exocrine lineages (11, 12). To date, a systematic exploration of DNA methylation changes during human pancreas development has not been performed due to, at least in part, the paucity of tissue from fetal donors. Several studies have utilized human fetal pancreatic samples to characterize transcription factor expression (1) and single cell gene expression patterns (13, 14) in the developing human pancreas, although these studies have focused on very early developmental stages and profiled only a limited number of samples. Our understanding of the genomic changes taking place during human pancreas development has therefore relied largely on extrapolations from animal model studies (15) and pancreatic cell lines (16, 17).

In this study we quantified DNA methylation across the genome in pancreas samples obtained from 99 fetal donors spanning 6 to 21 post-conception weeks (PCW) and 23 postnatal (adult) donors. We report dramatic changes in DNA methylation across human pancreas development, which are specific to the fetal period and distinct to age-associated changes in the postnatal pancreas. This is, to our knowledge, the most extensive study of DNA methylation across human fetal pancreas development and confirms the prenatal period as a time of considerable epigenomic plasticity.

## RESULTS

### Generating a high-quality DNA methylation dataset across human fetal pancreas development

We obtained flash-frozen 99 human fetal pancreas samples (43 male and 56 female) dissected from donors aged between 6 PCW (Carnegie stage (CS) 18) and 21 PCW from the Human Developmental Biology Resource (HDBR) (https://www.hdbr.org/) (**Figure 1A** and **Supplementary Table 1**). As expected, mean pancreas mass increased with age (**Supplementary** Figure 1) and immunohistochemical analysis of matched formalin-fixed paraffin embedded (FFPE) samples from a subset of donors (n = 10, aged 9 to 19 PCW, see **Methods**) was used to confirm the pancreatic origin of tissue from the HDBR prior to methylomic profiling (see **Figure 1B** and **Figure 1C**). We quantified DNA methylation across the genome in DNA isolated from each tissue sample using the Illumina Infinium MethylationEPIC (v1) array followed by a stringent quality control pre-processing pipeline (see **Methods**). After excluding poorly performing probes (i.e. those characterized by non-specific binding and those affected by genetic variation), in addition to those annotated to the Y chromosome, our final dataset included measurements of DNA methylation at 805,481 sites (787,690 (97.79%) autosomal, 17,791 (2.21%) on the X-chromosome)). We calculated a ‘DNA methylation age’ estimate for each sample using an epigenetic clock previously trained on fetal brain tissue and shown to be robustly associated with gestational age (18), finding a strong positive correlation between estimated age and actual developmental age (corr = 0.89, P = 8.31 x 10^-35^, **Figure 1D**). Finally, global levels of DNA methylation in the fetal pancreas, calculated by taking the mean across all autosomal DNA methylation sites included in the final dataset, did not differ significantly over the course of development (corr = 0.069, P = 0.497) (**Supplementary** Figure 2) indicating no overall changes in DNA methylation content across the genome during this period.

**FIGURE 1:**
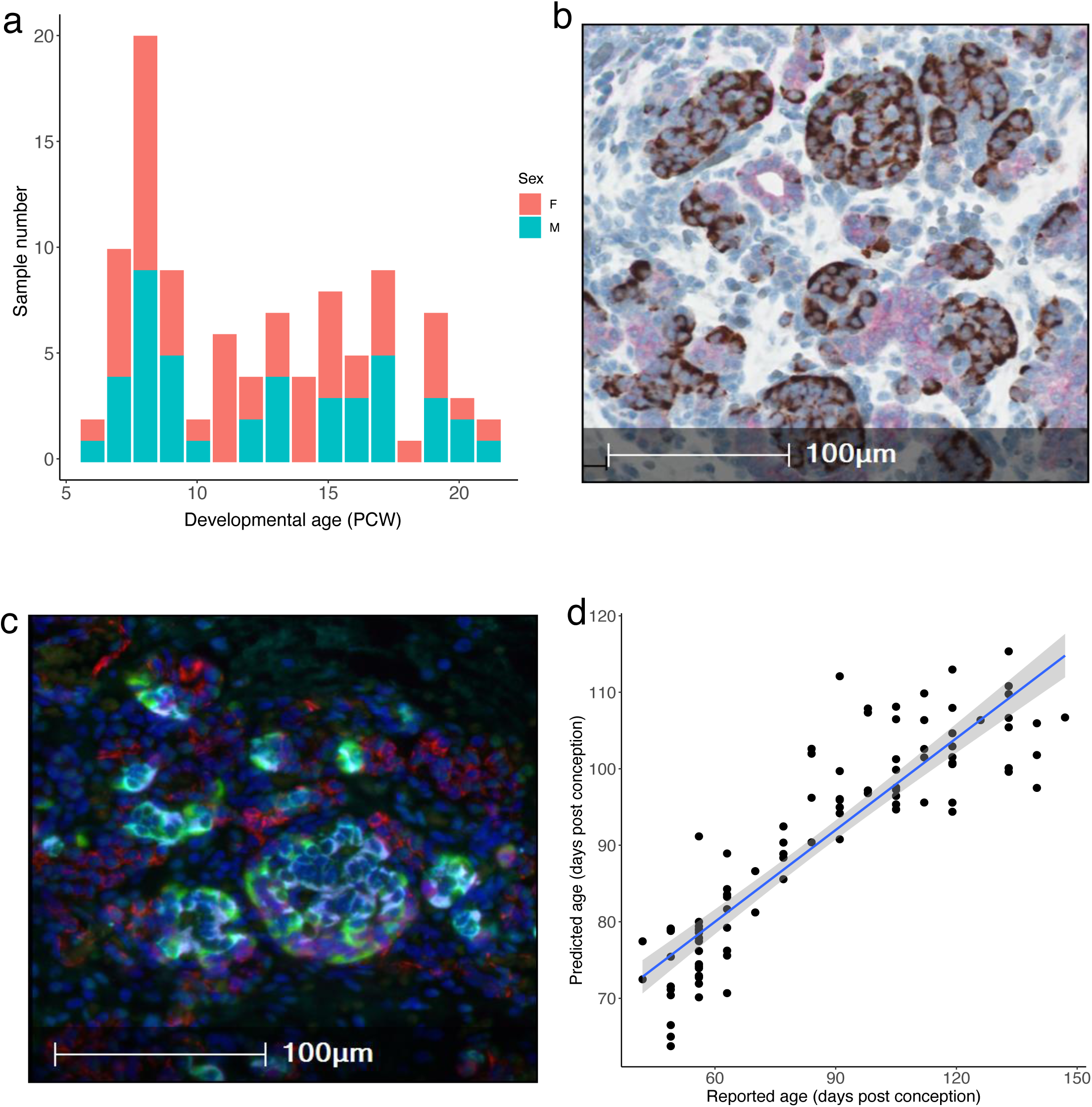
Characterization of fetal pancreas samples. **A)** An overview of the age and sex distribution of fetal pancreas samples profiled in this study. **B)** Chromogranin A (brown) and Cytokeratin 19 (pink) staining of FFPE tissue from a 17 PCW donor showing both endocrine and ductal tissue. **C)** Immunofluorescent staining analysis of FFPE tissue from a 17 PCW donor confirming the presence of endothelial cells (CD31+, red), alpha cells (GLU+, green), beta cells (INS+, cyan), and delta cells (SST+, magenta). **D)** Age estimates derived from an epigenetic clock calibrated on fetal tissue samples confirm a strong correlation between actual and estimated gestational age across the 99 fetal pancreas samples included in this study (corr = 0.89, P = 8.31 x 10^-35^).

### Development of the human fetal pancreas is associated with dramatic changes in DNA methylation at specific sites across the genome

We used a linear regression model, controlling for sex, to identify sites at which DNA methylation changed significantly across fetal pancreas development (see **Methods**); we refer to these as developmental differentially methylated positions (dDMPs). In total, we identified 177,130 dDMPs associated with developmental age at an experiment-wide significance threshold (*P* = 9 x 10^-8^), of which 174,992 (98.79%) were autosomal and 2,138 (1.21%) were located on the X chromosome (**Figure 2A** and **Supplementary Table 2**). Across all autosomal dDMPs there was a modest excess of sites characterized by decreasing levels of DNA methylation (‘hypomethylated dDMPs’) across fetal pancreas development (hypomethylated dDMPs = 88,345 (50.5%), hypermethylated dDMPs = 86,647 (49.5%), P = 9 x 10^-8^) (**Supplementary** Figure 3), consistent with patterns previously identified in our analyses of the developing human fetal brain (19).

**FIGURE 2:**
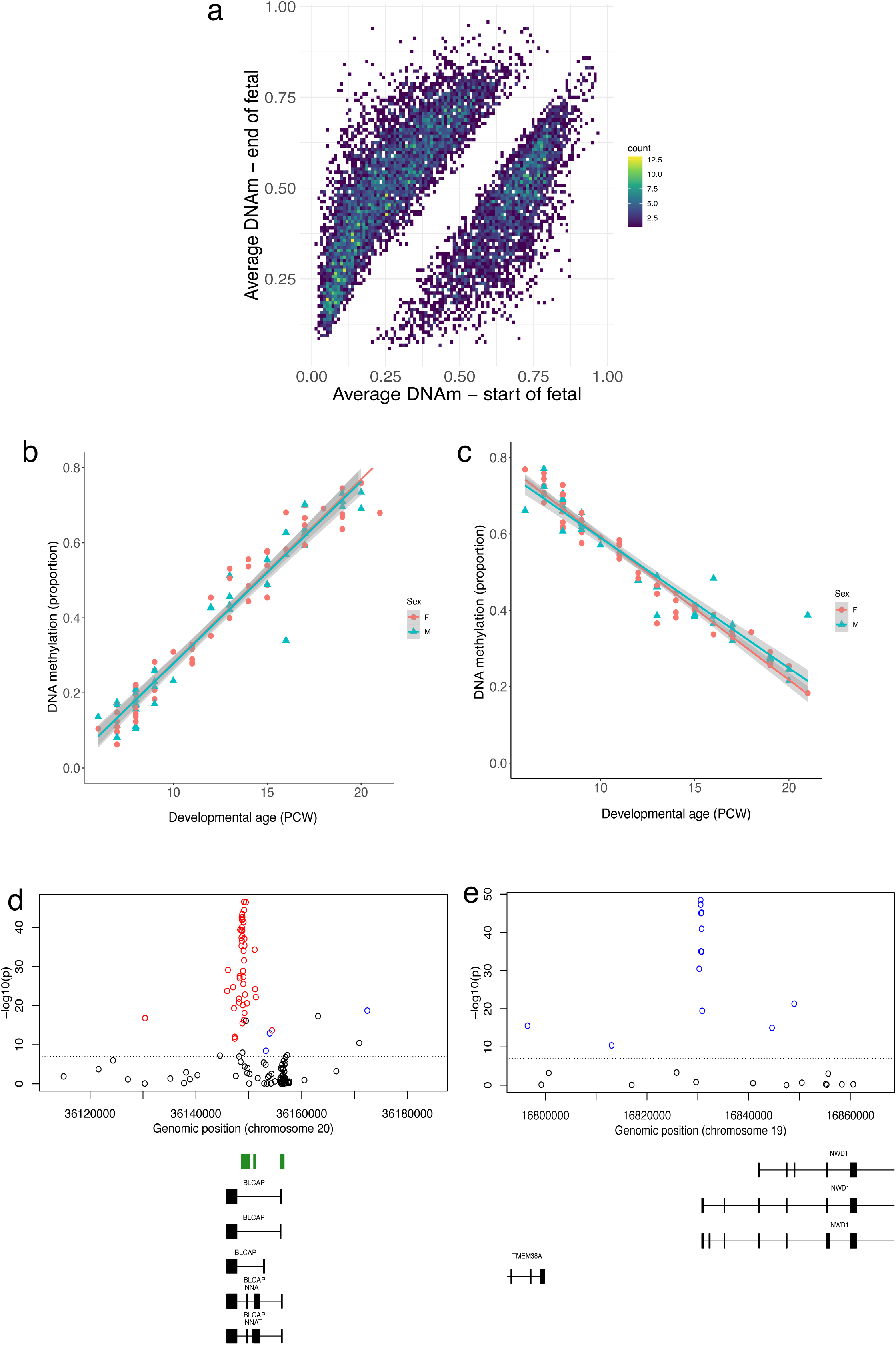
Changes in DNA methylation associated with development of the human pancreas. **A)** Mean levels of DNA methylation in early fetal pancreas samples (6 - 8 PCW) and later fetal pancreas samples (19 - 21 PCW) across all significant dDMPs. **B)** cg08125539, annotated to the *IGF2BP1* gene on chromosome 17, was the most significant hypermethylated dDMP across pancreas development (change in DNA mmethylation proportion per gestational week = 0.049, P = 1.35 x 10^-61^). **C)** cg20554008, annotated to the *MAP2K3* gene on chromosome 17, was the most significant hypomethylated dDMP across pancreas development (change in DNA methylation proportion per gestational week = - 0.036, P = 1.48 x 10^-61^). Many dDMPs are colocalized into larger regions of differential DNA methylation (dDMRs) associated with pancreas development. Shown are **D)** the top-ranked hypermethylated dDMR located in the *BLCAP* and *NNAT* genes on chromosome 20 (regression coefficient = 0.253, P = < 1.14 x 10^-304^) and the top-ranked hypomethylated dDMR located in the transcription start site of NWD1 on chromosome 19 (regression coefficient = -0.239, P = P = < 1.14 x 10^-304).^

The top-ranked dDMP associated with fetal pancreas development was cg08125539, located in intron two of the gene encoding insulin-like growth factor 2 mRNA binding protein 1 (*IGF2BP1*), which became progressively methylated over time (mean change in proportion DNA methylation per PCW = 0.049, P = 1.35 x 10^-61^, **Figure 2B**). *IGFBP1* is the predominant insulin-like growth factor (IGF)-1 binding protein during fetal development and is primarily secreted by the liver suggesting a role in the regulation of fetal growth (20). Interestingly, in adulthood, low IGFBP-1 levels are associated with a higher risk for type 2 diabetes (21). The top-ranked hypomethylated dDMP across pancreas development was cg20554008, located in the first intron of the gene encoding mitogen-activated protein kinase 3 (*MAP2K3*) (mean change in proportion DNA methylation per PCW = -0.036, P = 1.48 x 10^-61^, **Figure 2C**). *MAP2K3* is involved in the MAP kinase signaling pathways which has a critical role in cellular proliferation and differentiation in various contexts including cancer (22).

We performed pathway analysis on genes annotated to dDMPs using the *missMethyl* R package (23). Gene ontology (GO) analysis of genes annotated to the top 10,000 hypermethylated dDMPs identified a highly significant enrichment for multiple developmental pathways (**Supplementary Table 3**). KEGG analysis revealed an enrichment of genes within the Wnt signaling pathway (FDR = 0.0011), a crucial pathway for cell fate determination and organogenesis that plays an important role in development of the pancreas (24–26), and insulin secretion (FDR = 0.088) **(Supplementary Table 4)**. GO analysis of genes annotated to the top 10,000 hypomethylated dDMPs revealed an enrichment of genes involved in catalytic activity (FDR = 6.87 x 10^-8^) and other catabolic processes (FDR = 3.12 x 10^-7^) (**Supplementary Table 5**). KEGG analysis of hypomethylated dDMPs highlighting an enrichment of genes involved in endocytosis (FDR = 0.077) and protein processing in the endoplasmic reticulum (FDR = 0.077), both of which are implicated in the regulation of secretion from pancreatic β-cells (27) **(Supplementary Table 6**).

Many of the individual DMPs associated with pancreas development are in close proximity to each other, clustering into developmentally regulated differentially methylated regions (dDMRs). We used *dmrff* (28) to formally define these spatially-correlated regions of variable DNA methylation (see **Methods**). In total we identified 7,934 dDMRs (corrected P < 0.05, number of probes ≥ 3) spanning an average of 389 bp (with a range of 3-37 sites) (**Supplementary Table 7**). Many of the top-ranked dDMRs are proximal to genes with established roles in development and function of the pancreas. For example, the top hypermethylated dDMR (chr20: 36148264-36149656, 30 sites, *P* < 1.14 x 10^-304^, **Figure 2D**) spans the transcription start-site of the neuronatin gene (*NNAT*) that itself is located within the bladder cancer associated protein gene (*BLCAP*). *NNAT* is an imprinted proteolipid-encoding gene expressed specifically from the paternal allele that is highly expressed within differentiating endocrine cells and immature aggregating islet cells in mice (29, 30). Of note, a recent study showed that the mouse *Nnat* gene is differentially expressed across development in a discrete population of beta cells, and that this transcriptional plasticity is directly regulated by DNA methylation (31). The top hypomethylated dDMR (chr19: 16830287-16830859, 9 sites, *P* = < 1.14 x 10^-304^, **Figure 2E**) spans the transcription start site (TSS) of the gene encoding WD repeat domain-containing protein 1 (*NWD1*), which plays an important role in the developmental regulation of gene expression (32).

### The developing human pancreas has distinct modules of co-methylated loci

We used weighted gene correlation network analysis (WGCNA) (33) to undertake a systems-level characterization of the DNA methylation changes associated with human pancreas development. To enable a focus on sites annotated to genes we selected sites located in ‘promoter’ and ‘gene body’ regions characterized by variable DNA methylation, identifying discrete modules of co-methylated positions and used the first principal component of each individual module (the “module eigengene” (ME)) to assess their relationship with fetal pancreas development (see **Methods**). Among promoter-annotated sites (n = 92,989 variable sites) we identified 17 co-methylation modules of which 12 were significantly associated with pancreas development (Bonferroni-corrected *P* < 0.003) (**Supplementary Table 8**). The Promoter3 module (n = 5,091 probes) was the most positively correlated with developmental age (correlation of ME with age = 0.92, *P* = 2 x 10^-42^, **Figure 3A**) and the Promoter2 module (n = 5,955 probes) was characterized by the strongest negative correlation with developmental age (correlation of ME with age = -0.88, *P* = 2 x 10^-32^, **Figure 3B**). Module membership for sites included in both modules was found to be highly correlated with site significance with developmental age (Promoter3: r = 0.92, *P* < 1 x 10^-200^; Promoter2: r = 0.81, *P* < 1 x 10^-200^) (**Supplementary** Figure 4) indicating a clear relationship between DNA methylation at sites within these modules and fetal pancreatic development. Among gene body annotated sites (n = 77,945 variable sites), we identified 16 co-methylation modules, of which 13 were significantly associated with pancreas development (Bonferroni-corrected P < 0.003) (**Supplementary Table 9**). The GeneBody3 module (n = 10,312 probes) was most positively correlated with developmental age (correlation of ME with age = 0.919, *P* = 6.26 x 10^-41^, **Figure 3C**) and the GeneBody4 module (n = 8,419 probes) was most negatively correlated with developmental age (correlation of module eigengene (ME) = -0.741, *P* = 1.76 x 10^-18^, **Figure 3D**). Again, module membership for both modules was found to be highly correlated with site significance (GeneBody2 module: r = 0.88, P < 1 x 10^-200^; GeneBody3 module: r = 0.61, *P* < 1 x 10^-200^ (**Supplementary** Figure 5) indicating a clear relationship between DNA methylation at sites within these modules and fetal pancreatic development. To investigate the biological significance of the genes annotated to the core DNA methylation sites in each of the modules associated with pancreas development we conducted GO and KEGG pathway analysis (see **Methods**) identifying many relevant pathways including several related to the metabolism of macromolecules in the Promoter1 module (FDR = 0.001), bile secretion in the Promoter4 module (FDR = 0.008) and anatomical structure morphogenesis in the Promoter3 module (FDR = 0.02) (**Supplementary Table 10**).

**FIGURE 3:**
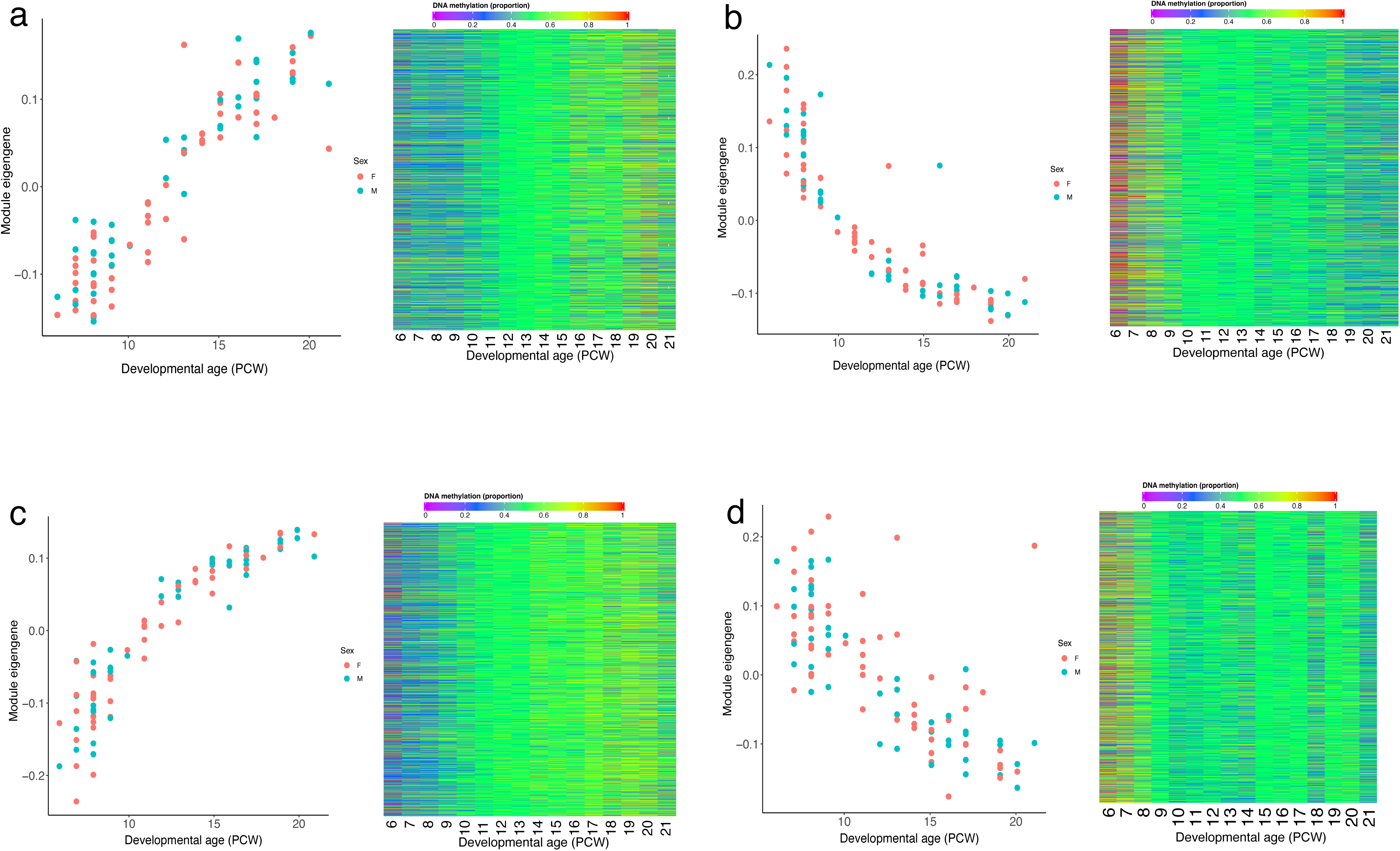
Gene comethylation modules associated with development of the human pancreas. Across DNA methylation sites located in promoter regions, the module eigengene of the **A**) Promoter4 module was most positively correlated with developmental age and the module eigengene of the **B)** Promoter3 module had the strongest negative correlation with developmental age. Shown for each module is the module eigengene for each sample (left panel) and a heatmap depicting the mean DNA methylation for core sites in each module across development (right panel) with colour corresponding to DNA methylation level at sites with a module membership >0.85. Across DNA methylation sites located in gene body regions, the module eigengene of the **C)** GeneBody3 module was was most positively correlated with developmental age and the module eigengene of the **D)** GeneBody4 module had the strongest negative correlation with developmental age.

### Autosomal sex-specific differences in DNA methylation in the developing pancreas

DNA methylation at 12,952 sites (1.61% of tested sites) was significantly different (*P* < 9 x 10^-8^) between males and females in the fetal pancreas. As expected, the vast majority of these differences occurred at sites located on the X chromosome (n = 12,242 (94.5%), mean male-female difference = 0.18 (SD = 0.12), range of male-female differences = 0.69 – 0.01), although a notable proportion were autosomal (n = 646 (4.99%), mean male-female difference = 0.048 (SD = 0.037), range of male-female differences = 0.30 – 0.009) (**Supplementary Table 11**). This may indicate sex-specific differences in gene regulation in the developing pancreas and is an interesting observation given evidence for male-female differences in pancreatic size and function (34, 35). The top-ranked autosomal DMP between males and females was cg06513015 (difference in mean DNA methylation = 0.27, *P* = 2.34 x 10^-78^) which is annotated to the promoter regions of the *ERV3-1* gene; of note this site has previously been shown to have significantly higher DNA methylation in newborn females compared to males (36, 37) (**Supplementary** Figure 6). Interestingly, we also identified 115 sites (all on the X chromosome) characterized by significant sex-specific changes in DNA methylation across development of the pancreas, potentially reflecting temporal changes in X chromosome dosage compensation mechanisms across development (**Supplementary Table 12** and **Supplementary** Figure 7).

### The distribution of dDMPs differs across genic features and is depleted in key transcription factor binding domains and regions of open chromatin

Although the distribution of dDMPs was largely consistent across autosomes (**Supplementary** Figure 8), certain genic features were highly enriched or depleted for developmentally-dynamic sites. For example, we observed a striking depletion of dDMPs in CpG islands (odds ratio = 0.241, P < 2.23 x 10^-308^), transcription start sites (TSS200) (odds ratio = 0.377, P < 2.23 x 10^-308^) and 1st exons (odds ratio = 0.389, P < 2.23 x 10^-308^) (**Figure 4A**). In contrast, there was a significant enrichment of dDMPs in features including CpG island shores (odds ratio = 1.10, P = 5.38 x 10^-52^), CpG island shelves (odds ratio = 1.14, P = 2.97 x 10^-39^), and gene bodies (odds ratio = 1.18, P = 3.63 x 10^-210^) (**Supplementary Table 13**). The depletion of dDMPs in CpG-rich promoter regions was driven primarily by a lack of sites becoming hypomethylated across development (**Figure 4B**); in contrast to other genic features we see a highly significant enrichment of hypermethylated dDMPs relative to hypomethylated dDMPs in CpG islands (sites characterized by decreasing DNA methylation = 4,294 (37.1%), sites characterized by increasing DNA methylation = 7,266 (62.9%), P < 2.2 x 10^-16^, **Supplementary** Figure 3). Reflecting this, mean DNA methylation across all dDMPs located in CpG islands significantly increases across development of the pancreas (mean change in DNA methylation per PCW = 0.0046, P < 2.2 x 10^-16^) (**Supplementary** Figure 2). These patterns reflect the fact that CpG islands are generally characterized by low levels of DNA methylation early in development with tissue-specific patterns of DNA methylation becoming established during the prenatal period (38).

**FIGURE 4:**
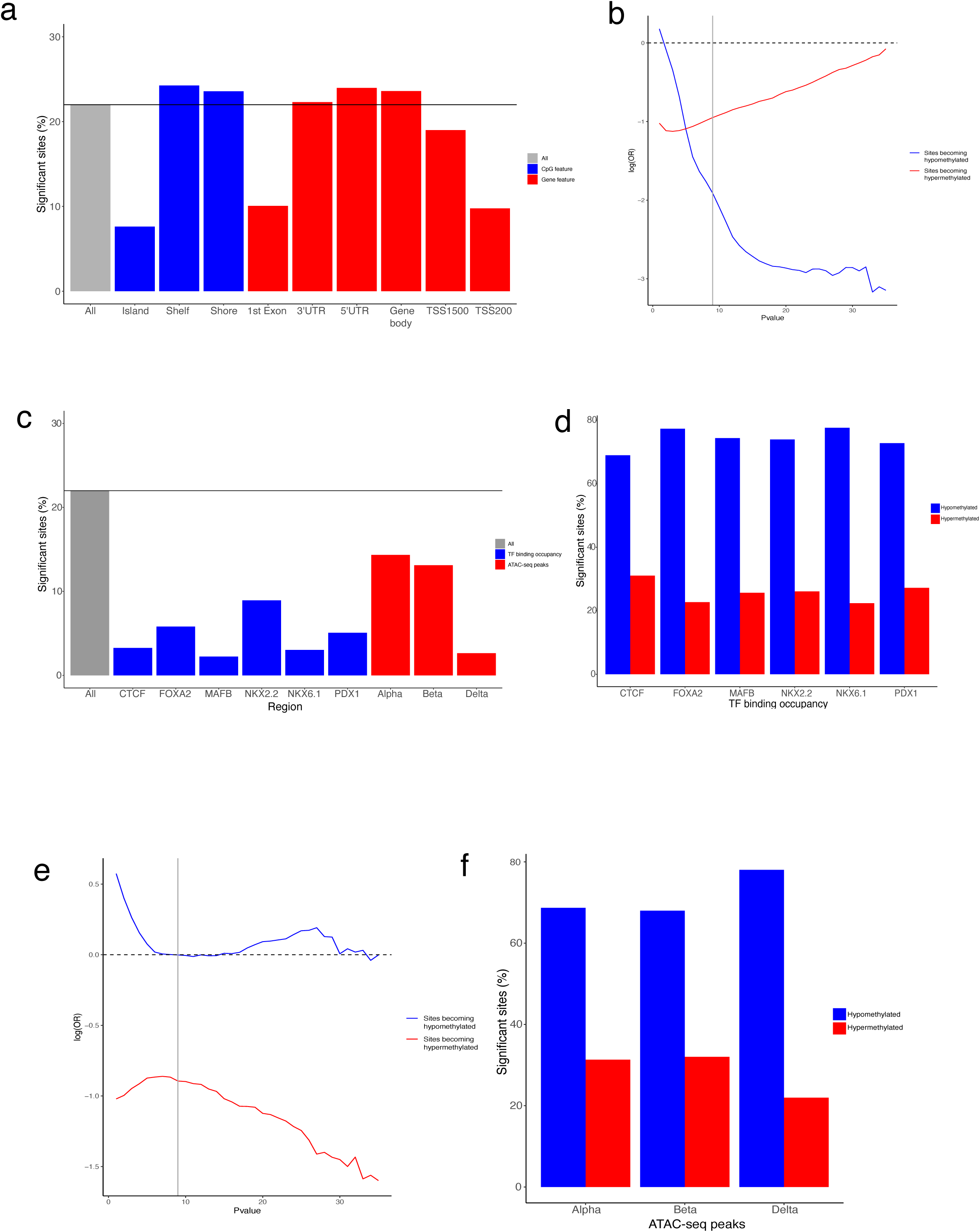
The distribution and direction of fetal pancreas dDMPs differs markedly across genomic features. **A)** Compared to the genome average, dDMPs are significantly underrepresented in CpG islands, transcription start-sites, 5′ UTRs, and first exons, but significantly enriched in CpG island shores, CpG island shelves, and the gene bodyRelative enrichment of dDMPs in CpG island features. **B)** The depletion of dDMPs in CpG islands is largely driven by a dramatic paucity of sites becoming hypomethylated across development. Sites becoming hypomethylated across development are shown in blue and sites becoming hypermethylated across development are shown in red. Shown is the relative enrichment of sites in each category based on their P-value for association with developmental age (from 1 x 10^-1^, to 1 x 10^-35^) determined using a Fisher’s exact test. The genome-wide significance threshold for dDMPs (P < 9 x 10^-8^) is denoted using a grey line. **C)** Compared to the genome average, dDMPs are also depleted in regulatory domains defined by key transcription factor binding site occupancy and chromatin accessibility in pancreatic islets from human donors. In contrast to the pattern seen in CpG islands, these regulatory domains are characterized by a dramatic depletion of hypermethylated dDMPs for both regions of **D)** transcription factor binding occupancy and **E)** open chromatin in pancreatic islet cell-types. **F)** The depletion of dDMPs in domains bound by CTCF, for example, is largely driven by a dramatic paucity of sites becoming hypomethylated across development.

We next sought to investigate whether dDMPs were enriched within known pancreatic regulatory domains identified from recent published analyses of key transcription factor binding site occupancy and chromatin accessibility in pancreatic islets from human donors. First, we used pancreatic islet ChIP-seq data generated on pancreatic islet cells (39) to test for an enrichment of dDMPs in peaks highlighting binding of the transcriptional regulator CCCTC binding factor (CTCF) and five pancreatic transcription factors (FOXA2, MAFB, NKX2-2, NKX6-1, and PDX1) (40–44). Mirroring the patterns observed in CpG islands, we found a highly significant depletion of dDMPs in domains bound by each of these transcription factors (**Figure 4C** and **Supplementary Table 13**). In contrast to CpG islands, however, transcription factor ChIP-seq peaks were dramatically depleted for hypermethylated dDMPs compared to background rates across the genome (**Figure 4D**) (CTCF: odds ratio = 0.423, P < 2.2 x 10^-308^; FOXA2: odds ratio = 0.459, P < 2.2 x 10^-308^; MAFB: odds ratio = 0.635, P = 1.78 x 10^-48^; NKX2-2: odds ratio = 0.334, P < 2.2 x 10^-308^; NKX6-1: odds ratio = 0.254, P < 2.2 x 10^-308^; PDX1: odds ratio = 0.399, P = < 2.2 x 10^-308^) (**Figure 4E** and **Supplementary** Figure 9), reflecting the observation that the binding of many transcription factors is sensitive to methylated DNA (45, 46). Transcription factor binding occurs mainly in regions of open chromatin. We therefore used publicly available single nucleus ATAC-seq data to test the enrichment of dDMPs in genomic regions of open chromatin in three endocrine cell types (alpha-, beta-and delta-cells) (47). In all three cell types, regions of open chromatin were again characterized by an overall depletion of dDMPs driven by a highly significant depletion of hypermethylated dDMPs, mirroring the patterns seen in occupied transcription-factor binding sites (**Figure 4F** and **Supplementary** Figure 10). This depletion was particularly strong in delta cells where there was also an overall depletion of hypomethylated dDMPs (all dDMPs: odds ratio = 0.221, P < 2.2 x 10^-308^; hypermethylated dDMPs: odds ratio = 0.106, P < 2.2 x 10^-308^; hypomethylated dDMPs: odds ratio = 0.394, P < 2.2 x 10^-308^).

### A number of neonatal diabetes genes are enriched for dDMPs

Maturity onset diabetes of the young (MODY) and neonatal diabetes are monogenic disorders of the pancreas caused by pathogenic mutations in several genes, including those encoding many of the key transcription factors involved in coordinating pancreas development (48). We explored whether DNA methylation sites annotated to both MODY (N = 33, **Supplementary Table 14**) (https://panelapp.genomicsengland.co.uk/panels/472/) and neonatal diabetes (N = 32, **Supplementary Table 15**) (https://panelapp.genomicsengland.co.uk/panels/293/) genes (**Supplementary** Figure 11) were enriched for dDMPs associated with pancreas development. There was evidence for dDMPs annotated to all the tested MODY genes (33 genes (100%)) and virtually all tested neonatal diabetes genes (30 genes (93.7%)) genes (**Supplementary Table 14&15**). Although there was no overall significant enrichment of dDMPs to either MODY (odds ratio = 1.148, P = 0.021) or neonatal diabetes (odds ratio = 1.084, P = 0.133) genes, there was a significant enrichment of sites becoming hypermethylated across development annotated to neonatal diabetes gene regions (odds ratio = 1.357, P = 2.98 x 10^-4^) (**Figure 5A** and **Supplementary** Figure 12). A number of disease genes were characterized by very dramatic changes in DNA methylation across development. The neonatal diabetes gene with the highest proportion of significant dDMPs was *NKX2-2* encoding homeobox protein NKX2-2, a crucial islet transcription factor (42, 49). DNA methylation at 16 of 22 (72.72%) *NKX2-2* sites changed significantly across development (**Figure 5B**), with these sites clustered into a discrete dDMR spanning the gene characterized by increasing DNA methylation (beta = 0.195, P = 1.14 x 10^-51^) (**Figure 5C** and **Supplementary Table 7**). Another neonatal diabetes gene containing a dDMR characterized by dramatic hypermethylation across pancreas development encodes the lipopolysaccharide responsive beige like anchoring protein (*LRBA*); DNA methylation at 44 of 118 (37.29%) *LRBA* sites was associated with developmental age (**Figure 5D**), with these sites clustered into a discrete dDMR encompassing an intragenic CpG island (beta = 0.235, P = 1.14 x 10^-301^) (**Figure 5E** and **Supplementary Table 7**). Summary results for all neonatal diabetes genes are presented in **Supplementary Table 16**, potentially aiding the identification of regulatory domains with a critical role in pancreas development.

**FIGURE 5:**
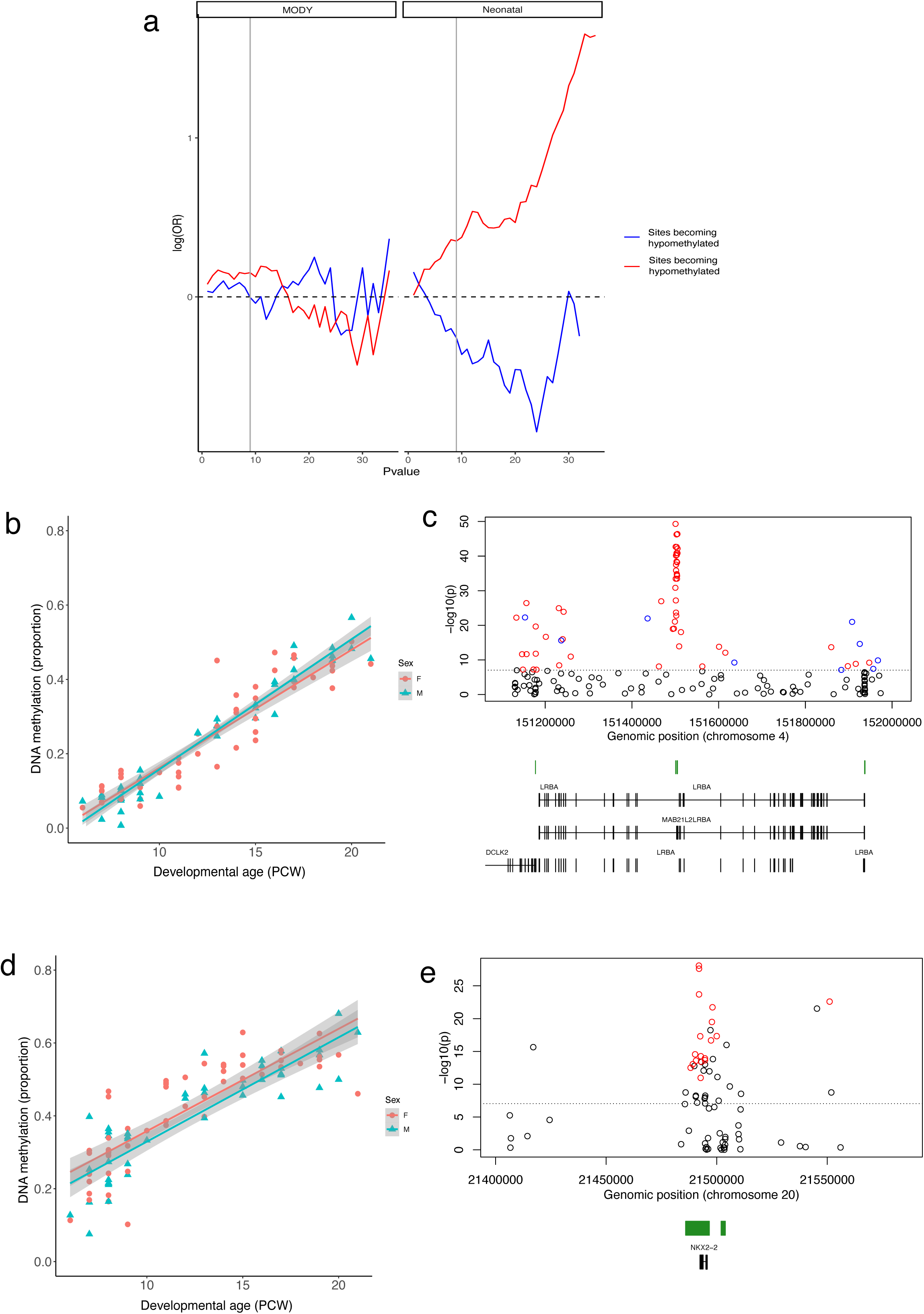
Developmental changes in pancreatic DNA methylation annotated to neonatal diabetes genes. **A)** There is a significant enrichment of sites becoming hypermethylated across pancreas development annotated to neonatal diabetes genes. Sites becoming hypomethylated across development are shown in blue and sites becoming hypermethylated across development are shown in red. Shown is the relative enrichment of sites in each category based on their P-value for association with developmental age (from 1 x 10^-1^, to 1 x 10^-35^) determined using a Fisher’s exact test. The genome-wide significance threshold for dDMPs (P < 9 x 10^-8^) is denoted using a grey line. **B)** cg23425348 shows the most significant developmental change in DNA methylation associated across sites annotated to *NKX2-2* (change in DNA methylation proportion per gestational week = 2.83 x 10^-2^, P = 8.26 x 10^-29^) which is the neonatal gene with the highest proportion of dDMPs and **C)** contains a discrete region of significant hypermethylation (regression coefficient = 0.195, P = 1.14 x 10^-51^). **D)** cg25995955 shows the most significant developmental change in DNA methylation associated across sites annotated to *LRBA* (change in DNA methylation proportion per gestational week = 3.33 x 10^-2^, P = 5.05 x 10^-50^). **F)** The LRBA gene is another neonatal diabetes gene with an enrichment of dDMPs clustered into a discrete region characterized by increasing DNA methylation across development (regression coefficient = 0.224, P = 1.14 x 10^-304^).

### The dramatic temporal changes in DNA methylation occurring in the developing pancreas are limited to the prenatal period and are distinct to age-associated changes in the postnatal pancreas

Given the notable shifts in DNA methylation observed across pancreatic development, we were interested in characterizing the extent to which these changes continued in the postnatal pancreas. We quantified DNA methylation across the genome in 23 adult pancreas samples obtained from the Barts Pancreas Tissue Bank (BPTB) (https://www.bartspancreastissuebank.org.uk/) (mean age = 59 years, age range 34 to 80 years, **Supplementary Table 1**) using the Illumina EPIC array. As expected, the chronological age of postnatal donors was strongly correlated with epigenetic age estimates derived from our DNA methylation data using the Horvath multi-tissue epigenetic clock (50) (corr = 0.88, P = 2.91 x 10^-8^, **Supplementary** Figure 13). We next tested the extent to which dDMPs were also characterized by age-associated changes in DNA methylation in postnatal pancreas. Across the 176,624 significant fetal pancreas dDMPs also tested in the adult pancreas, there was no positive correlation in effect sizes (**Figure 6A**), suggesting that most of the dramatic temporal changes at fetal dDMPs is limited to the prenatal period and do not reflect aging *per se*. Overall, DNA methylation at pancreas dDMPs was more variable in fetal pancreas samples than postnatal pancreas samples (**Figure 6B**) (average variance in DNA methylation across the top 10,000 dDMPs profiled in both datasets = 0.0150 (fetal pancreas) vs 0.0013 (postnatal pancreas), P = 9 x 10^-8^). DNA methylation at pancreas dDMPs was markedly different between early fetal (6 - 8 PCW) and adult pancreas samples (corr = -0.206, **Figure 6C**), mean levels of DNA methylation in adult pancreas (average age = 59 years) were strongly correlated (corr = 0.697) to those seen older fetal samples (19-21 PCW). These results further highlight the relative stability of DNA methylation at these sites postnatally following the dramatic shifts observed during pancreatic development (**Figure 6D**).

**FIGURE 6:**
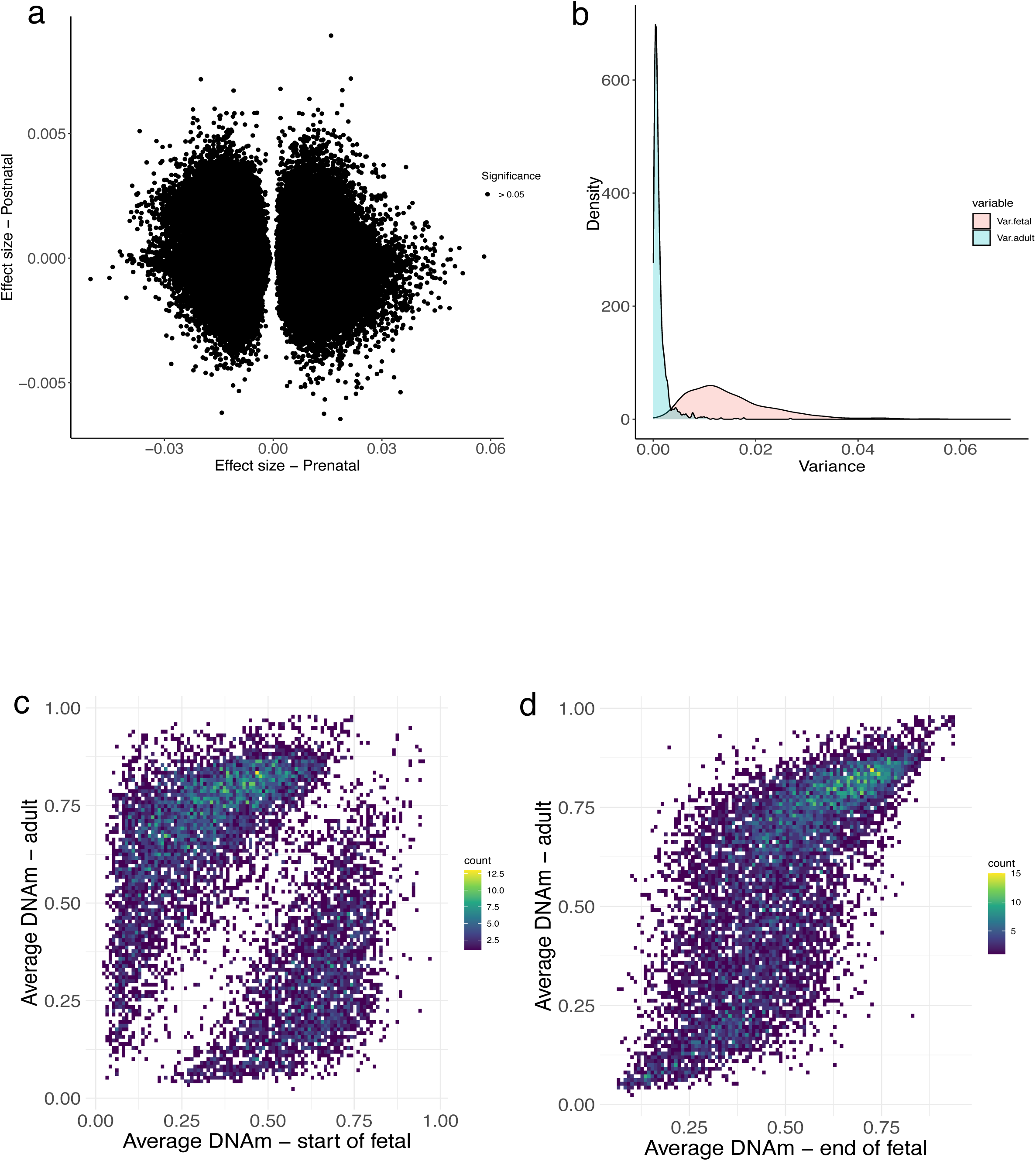
The dramatic temporal changes in DNA methylation at fetal dDMPs is limited to the prenatal period. **A**) Correlation of DNA methylation changes during fetal development (X-axis) and postnatal aging (Y-axis) at 176,624 dDMPs tested in both fetal and adult pancreas. No pancreas dDMPs were significantly associated with age in the postnatal pancreas. **B**) DNA methylation at pancreas dDMPs was dramatically more variable in fetal pancreas samples than postnatal pancreas samples. Shown is the distrinution of variance in DNA methylation across the 10,000 most significanty dDMPs in fetal pancreas samples (pink) (variance = 0.0150) compared to the variance in DNA methylation at the same sites in adult pancreas samples (blue) (0.0013). **C**) DNA methylation at pancreas dDMPs was dramatically different between early fetal (6 - 8 PCW) and adult pancreas samples at the 10,000 most significant dDMPs. **D**) In contrast, DNA methylation at pancreas dDMPs was highly correlated between early fetal (6 - 8 PCW) and adult pancreas samples at the same dDMPs (corr = 0.70).

### There are tissue-specific developmental changes in DNA methylation specific to the pancreas

Using data from our previous study of DNA methylation changes during development of the human brain (19) we explored the extent to which dDMPs identified in this study were specific to development of the pancreas. 73,964 (41.75%) of the identified pancreas dDMPs were also profiled in the human brain using an earlier version of the Illumina array (Illumina 450K); across these sites there was a modest positive correlation in effect size between pancreas and brain (corr = 0.277, P = < 2.2 x 10^-16^) (**Figure 7A** and **Supplementary Table 17**). DNA methylation at 8,131 (10.99%) of the tested pancreas dDMPs was significantly associated with development in the human brain at a Bonferroni-corrected significance threshold (P < 6.76 x 10^-7^). Of these, DNA methylation at 6,645 (81.72%) sites was characterized by a consistent direction of developmental change in pancreas and brain (e.g., **Figure 7B**) whilst 1,486 (18.28%) sites were characterized by an opposite direction of effect across tissues (e.g., **Figure 7C**). A relatively large number of DMPs were also found to be specific to the developing pancreas; in total we found 55,402 pancreas DMPs (74.90%) that did not appear to be dynamically regulated in the developing brain (i.e., pancreas DMPs at which there was a non-significant mean change in proportion DNA methylation per PCW <0.005 and >-0.005 in the fetal brain) (**Supplementary Table 19**). Of note, the top-ranked example of a pancreas-specific DMP was annotated to the neonatal diabetes gene *LBRA* (cg08219218: absolute difference in mean change in DNA methylation per PCW between the fetal pancreas and brain = 0.0527, **Figure 7D**). We also identified pancreas-specific changes across sites annotated to *NNAT* within the top-ranked dDMR, suggesting that developmental changes in DNA methylation across this domain are specific to the pancreas (**Supplementary** Figure 14).

Similarly, of the 13,480 brain dDMPs identified by Spiers and colleagues at P < 9 x 10^-8^, 12,531 (92.96%) were included in our fetal pancreas dataset; there was an overall positive correlation in effect sizes across the two tissues (corr = 0.483, P = < 2.2 x 10^-16^) (**Figure 7E** and **Supplementary Table 18**). 6,831 (54.51%) of the tested brain dDMPs were significantly associated with developmental age in the human pancreas at a Bonferroni-corrected significance threshold (P < 3.99 x 10^-6^), with 5,513 (80.71%) sites characterized by a consistent direction of effect in the brain and 1,318 (19.29%) sites characterized by an opposite direction of effect in the brain. We identified a number of brain-specific dDMPs with 1,545 sites (12.33%) not characterized by developmental changes in the developing pancreas (**Supplementary Table 20**). Of note, the top-ranked brain DMP at which DNA methylation did not change during pancreas development was annotated to the Methionine Sulfoxide Reductase A gene (*MSRA*) (cg02192555: absolute difference between change in mean DNA methylation per PCW between the fetal brain and pancreas = 0.0368) (51). Overall, these results indicate that whilst many dDMPs show very similar developmental trajectories across tissues (**Figure 7C**), a relatively large proportion of sites are characterized by tissue-specific developmental changes in DNA methylation (**Figure 7D**).

**Figure 7:**
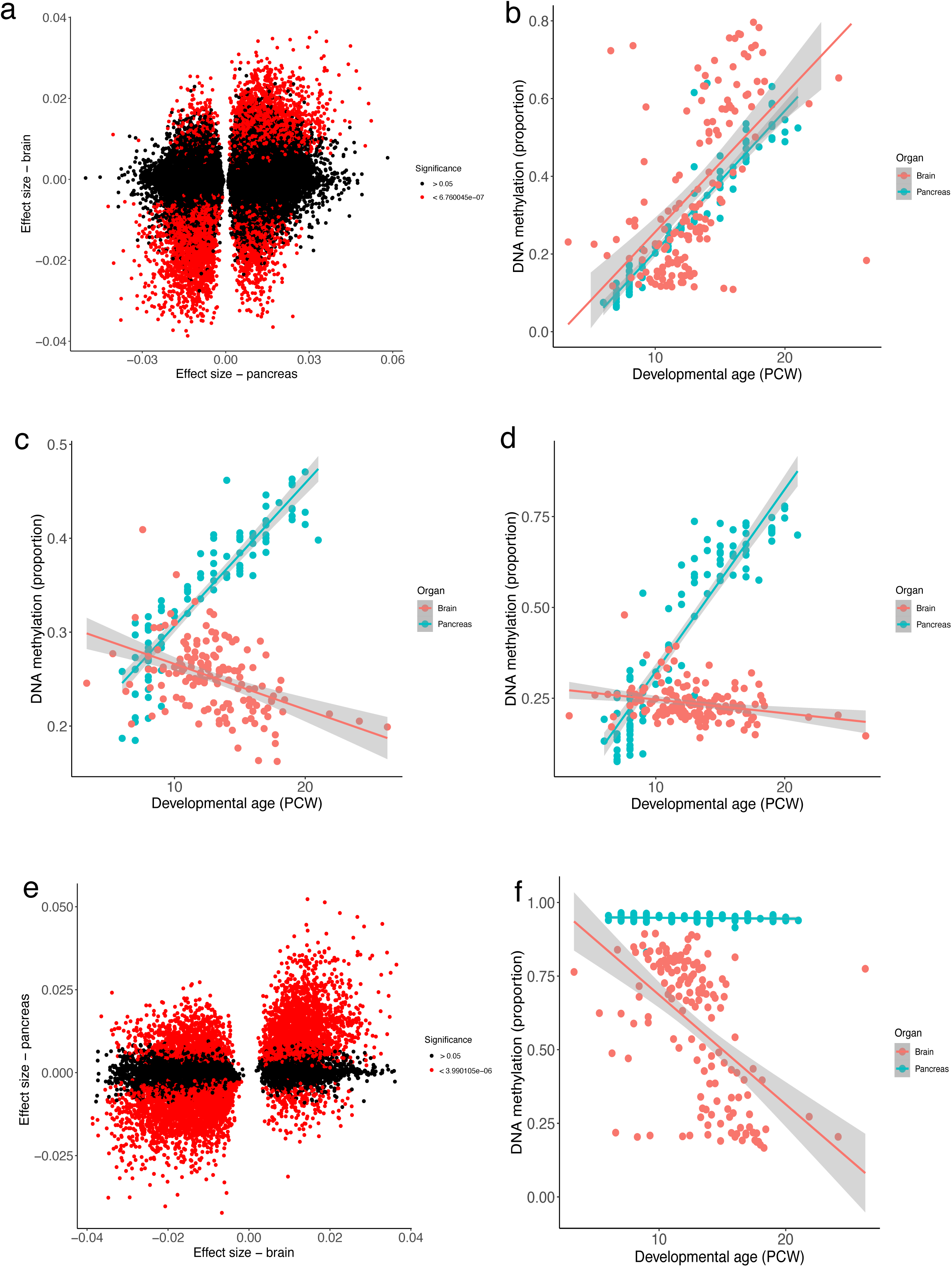
Overlap between developmental changes in DNA methylation between pancreas and brain. **A)** Correlation of effect sizes at pancreas dDMPs with those at the same sites in the developing brain. **B)** A site at which developmental changes in DNA methylation is highly similar between pancreas and brain. Shown is cg08486065 which is characterized by significant decrease in DNA methylation across development in both pancreas (effect size = 0.0362, P = 1.06 x 10^-54^) and brain (effect size = 0.0341, P = 1.38 10^-12^). **C)** A site at which DNA methylation changes significantly across development in the pancreas and brain but in the opposite direction. Shown is cg16094352 which is characterized by significant hypermethylation across development in the pancreas (effect size = 0.0151, P = 3.13 x 10^-41^) but significant hypomethylation across development in the brain (effect size = -0.00484, P = 1.68 x 10^-8^). **D)** A site at which developmental changes in DNA methylation are specific to the pancreas. Shown is cg08219218 annotated to *LBRA* that is characterized by significant hypermethylation across development in the pancreas (effect size = 0.0490, P = 4.52 x 10^-41^) but no change in the brain. **E)** Correlation of effect sizes at brain dDMPs with those at the same sites in the developing pancreas. **F)** A site at which developmental changes in DNA methylation are specific to the brain. Shown is cg02192555 annotated to *MSRA* that is characterized by significant hypomethylation across development in the brain (effect size = -0.0371, P = 3.86 x 10^-12^) but no change in the pancreas.

## DISCUSSION

In this study, we characterized changes in DNA methylation occurring through development of the human pancreas. We profiled genome-wide patterns of DNA methylation in pancreas samples from 99 fetal donors (43 male and 56 female) spanning 6 to 21 post conception weeks, and 23 postnatal (adult) pancreas samples (ages 34 to 80 years, 9 male and 14 female). Our data highlight that considerable epigenomic changes taking place in the human pancreas during fetal development, with significant changes in DNA methylation identified at over 21% of sites tested. The distribution of pancreas dDMPs was found to differ across genic regions, with notable depletion in CpG islands, regions of open chromatin and the binding sites of pancreatic transcription factors involved in pancreas development. The dramatic temporal changes in DNA methylation occurring in the developing pancreas were found to be limited to the prenatal period and be distinct to age-associated changes observed in the postnatal pancreas. Finally, we compared our data with those from a similar study on the developing human brain, finding both common developmental changes across tissues and tissue-specific trajectories of DNA methylation. This is, to our knowledge, the most extensive study of DNA methylation across human fetal pancreas development to date, confirming the prenatal period as a time of considerable epigenomic plasticity.

Distinct developmental patterns of DNA methylation were observed in several genomic regions. DNA methylation in CpG-rich areas were found to exhibit less variability during fetal pancreas development compared to other genomic regions. Moreover, although CpG-rich regions tend to have more hypermethylated dDMPs than hypomethylated dDMPs (**Supplementary** Figure 3), the majority of dDMPs in the genome become hypomethylated as the pancreas develops. This observation is unsurprising considering that CpG islands play a role in stabilizing the expression of essential housekeeping genes (6) and reflects our previous observations in the developing human brain (19). We also investigated changes in DNA methylation within regulatory regions bound by specific islet transcription factors important for pancreas development and function (i.e. FOXA2, NKX2-2, NKX6-1, PDX1, MAFB, and CTCF). DNA methylation is known to disrupt the developmental binding of many transcription factors (6), and therefore the progressive hypomethylation observed across these domains may orchestrate the complex patterns of gene expression crucial for pancreatic development and function. In particular, the binding of CTCF, which plays a crucial role in the developing pancreas by contributing to the establishment and regulation of chromatin architecture, gene expression and cell fate determination (52), is known to be highly anticorrelated with DNA methylation (53). These data support a critical role for developmental changes in DNA methylation during the differentiation of different pancreatic cell types, including those responsible for insulin production and glucose regulation.

We also investigated changes in DNA methylation across pancreas development in relation to sites annotated to genes associated with diseases of the pancreas including MODY and neonatal diabetes, finding that the vast majority of these genes contained at least one significant dDMP. Although MODY genes showed no enrichment for sites characterized exclusively by either increasing or decreasing DNA methylation across pancreatic development, there was a strong enrichment for sites to become hypermethylated in genomic regions annotated to neonatal diabetes genes. This is interesting given the role of aberrant pancreas development in the development of neonatal diabetes, a rare form of diabetes that presents in the first six months of life, and the fact that many of the genetic mutations that cause neonatal diabetes are known to disrupt normal pancreas development and function.

Our data may help elucidate novel mechanisms involved in diseases of the pancreas. One of the most significant dDMRs in our dataset, for example, was located within the *LRBA* gene (**Figure 5D**). Although *LRBA* mutations result in neonatal diabetes and additional autoimmune features diagnosed as early as six weeks of age (54). These mutations are thought to cause diabetes through de-regulation of the immune system and there is little previous evidence to suggest a role for *LRBA* in the development of the pancreas. Our data reveals that there are dynamic changes in the epigenetic state of LRBA across pancreas development, with progressive hypermethylation across a discrete region within the gene. Not all cases of neonatal diabetes and MODY can be explained by identified coding mutations in the coding region of known genes and understanding the epigenomic changes occurring during fetal pancreas development could provide clues to the location of novel (potentially non-coding) functional genetic variants contributing to the etiology of these disorders.

There are several limitations to this study. First, legal restrictions on later-term abortions precluded the assay of pancreas tissue from later stages of fetal development. However, the observation that DNA methylation at dDMPs was relatively stable between the oldest fetal samples studied and postnatal (adult) pancreas (**Figure 6D**) implies that the magnitude of epigenetic changes occurring later in pregnancy and during early postnatal life is much lower that was observed across early fetal development. Another limitation is that the tissue used in the present study was whole pancreas and our measurements of DNA methylation represent a composite of different cell types. This precludes us from exploring changes within specific cell-types in the developing pancreas, and future work should focus on profiling changes in purified cell-types. Third, although the Illumina EPIC array can accurately quantify DNA methylation at single-base resolution with probes associated with the majority of genes and CpG islands across the genome, the array targets only ∼2% of the CpG sites in the human genome, and probes are not equally distributed across all genomic features. As costs diminish, future studies should employ sequencing-based genomic-profiling technologies to more thoroughly interrogate the epigenome across development in large numbers of samples. Fourth, no demographic or phenotypic information other than developmental age and sex was available for the samples assayed. Fifth, we do not have gene expression data from these samples and are therefore unable to make direct conclusions about the transcriptional consequences of the developmental changes in DNA methylation we observe. Finally, we are not able to distinguish between DNA methylation and its oxidized derivative, DNA hydroxymethylation, as standard sodium bisulfite approaches do not distinguish between these modifications. There is currently very limited information about the prevalence of hydroxymethylated DNA in the pancreas and future studies should aim to characterize this modification across fetal pancreas development using techniques capable of discriminating DNA methylation and DNA hydroxymethylation (55–57).

In conclusion we identify highly dynamic changes in DNA methylation occurring throughout the genome across human fetal pancreatic development. The dramatic temporal changes in DNA methylation occurring in the developing pancreas appear to be limited to the prenatal period and be distinct to age-associated changes observed in the postnatal pancreas. The majority of genes involved in neonatal diabetes and MODY were located proximal to sites showing developmental changes in DNA methylation, potentially providing novel insights into the developmental mechanisms involved in these diseases.

## METHODS

### Human pancreas tissue samples

Human fetal tissues were obtained from the Human Developmental Biology Resource (HDBR). These samples were collected with appropriate maternal written consent and approval from the Newcastle and North Tyneside (Newcastle University) and London - Fulham (UCL) Research NHS Health Authority Joint Ethics Committees. HDBR is regulated by the UK Human Tissue Authority (HTA; www.hta.gov.uk) and operates in accordance with the relevant HTA Codes of Practice. The age of these samples ranged from 6 to 21 PCW, determined by Carnegie staging in the case of embryonic samples (defined as ≤56 DPC (8 PCW) and foot and knee to heel length measurements for fetal samples (defined as ≥57 DPC (8.14 PCW). Other than sex, no additional phenotypic or demographic information on the fetal donors was available. Human adult pancreatic tissue (n = 23) was acquired from the Barts Pancreas Tissue Bank at Queen Mary’s University London (www.bartspancreastissuebank.org.uk) with full approval from the Research Ethics Committee (REC) (references 13/SC/0592, 18/SC/0630 and 23/SC/0324). All samples used in this study were collected from the Barts Pancreas Tissue Bank, with written informed consent from patients recruited at Barts Health NHS Trust (BHNT). Pancreas samples were taken for surgical biopsy for pancreatic tumors, and only samples with no malignancies or cellular abnormalities were selected for this study.

### Immunohistochemistry (IHC) and multiparameter immunofluorescence imaging

2µm formalin-fixed paraffin-embedded (FFPE) pancreatic tissue sections obtained from HDBR were stained for the presence of chromogranin A (ChrgA) positive endocrine cells and CK19-positive ductal cells. Sections were double-stained with antibodies against ChrgA and CK19 using standard immunohistochemical approaches. Counterstaining with haematoxylin allowed for identification of exocrine and structural tissue. The source, characteristics, dilutions and validations of primary antibodies, as well as source, description and details of accessory agents are provided in **Supplementary Table 21**. For multiplex tissue immunofluorescence, 2µm FFPE pancreas tissue sections were baked at 60°C for 1 hour, dewaxed in histoclear, rehydrated in degrading ethanols (100%, 95%, 70%) and fixed in 10% neutral buffered formalin (NBF). Heat-induced epitope retrieval (HIER; 10mM citrate pH6) was performed to unmask the epitopes by placing the sections in a pressure cooker in a microwave oven on full power for 20 mins. The sections were then blocked with 5% normal goat serum (NGS) and incubated with the primary antibody before being probed with an appropriate OPAL™ secondary antibody conjugated to a fluorophore (Akoya Biosciences, **Supplementary Table 21**). This was followed by a further HIER to remove the primary antibody before staining with the next primary/secondary combination. The same steps (from blocking to antigen retrieval) are repeated four times (for each of the primary/secondary combinations). The sections are counterstained with DAPI and mounted for multispectral fluorescent microscopy performed using the PhenoImager HT automated quantitative pathology imaging system (Akoya Biosciences).

### Genome-wide quantification of DNA methylation and data preprocessing

Frozen pancreas tissue was homogenized using a motorized pestle prior to conducting nucleic acid extraction using the Qiagen AllPrep Universal kit (Qiagen). The processing of all samples was randomized to minimize batch effects attributed to age or sex. Genomic DNA (∼500ng) was treated with sodium bisulfite using the Zymo EZ DNA Methylation-Lightning kit (Zymo Research). DNA methylation was profiled across the genome using the Illumina Infinium MethylationEPIC BeadChip (Illumina) and scanned on the HiScan System (Illumina) using the standard manufacturers instructions. Data quality control and pre-processing were performed using the *wateRmelon* package as described previously (58). Cross-reactive probes and polymorphic DNA methylation sites, as detailed in the Illumina annotation file and identified in publications (59) were removed leaving 805,481 probes for further analysis (including n = 17,516 X chromosome and n = 77 Y chromosome probes) for fetal samples and 857,808 probes for adult pancreatic samples. Data were normalized with the *dasen* function of the *wateRmelon* package. Demographic information for the final cohort of 99 fetal pancreas samples and 23 adult pancreas samples remaining following quality control are provided in **Supplementary Table 1**.

### Statistical Analysis

A multiple linear regression model with developmental age (in post conception weeks, PCW) and sex was implemented to identify DNA methylation changes in the developing human pancreas. DNA methylation at specific sites was considered to be significantly associated with development (in PCW) and sex if they passed an experiment-wide significance threshold of P = 9 x 10^-8^. Analysis of differentially methylated probes at the regional level was performed using the DMR-finding tool *dmrff* (https://github.com/perishky/dmrff). Gene annotations provided by Illumina were used to map sites to specific genes. The *missMethyl* package (23) was used to conduct pathway analysis using standard parameters and relevant backgrounds.

### Enrichment analysis

We used publicly available ChIP-seq data generated on pancreatic islet cells (Pasquali 2014) to test for an enrichment of dDMPs in regions characterized by the binding of six transcription factors (CTCF, FOXA2, MAFB, NKX2-2, NKX6-1, and PDX1). We used publicly available single nucleus ATAC-seq data to test the enrichment of dDMPs in genomic regions of defined by open chromatin in three endocrine cell types (alpha-, beta-and delta-cells) (Chiou, *et al*. 2021). Diabetes gene lists were obtained from the PanelApp webpage of Neonatal diabetes genes and Monogenic diabetes genes. For analyses of different genic regions, we used feature annotations from the Illumina EPIC array manifest. For DNA methylation sites in each region, we calculated Fisher exact test P-values and odds ratios for the association between a site being present in a given set of genomic regions or genes and the site being differentially methylated with a P-value less than a given threshold (varying the threshold from P = 10^-1^ to P = 10^-35^).

### Weighted gene correlation network analysis (WGCNA)

Weighted gene correlation network analysis (WGCNA) was performed to identify modules of highly correlated DNA methylation sites associated with pancreas development (33). WGCNA was performed separately for DNA methylation sites annotated to promoter regions (n = 92,989 sites) and gene body regions (n = 77,945 sites). Modules were identified based on pairwise correlations using a signed network at power 4. We excluded modules containing <100 DNA methylation sites. A weighted average expression profile representing the first principal component of each individual module, otherwise known as a module eigengene (ME), was calculated for each module. Correlations between the MEs and the phenotypic traits available (PCW, sex) were used to identify modules associated with these traits. The relationship between each probe and each module was assessed through calculation of module membership (MM), the absolute correlation between a probe’s DNA methylation status and the ME, allowing the identification of the subset of probes with the strongest membership to each module. To assess the biological meaning of modules associated with pancreas development, genes associated with probes possessing an absolute MM value of >0.85 were extracted and assessed using pathway and gene ontology analysis.

## RAW DATA AVAILABILITY

Raw and normalized Illumina EPIC methylation data has been submitted to the NCBI Gene Expression Omnibus (GEO; http://www.ncbi.nlm.nih.gov/geo/) under accession number GSE244636.

## ACKNOWLEDGEMENTS

A.M. was supported by the UKRI Expanding Excellence in England award via EXCEED. Data analysis was undertaken using high-performance computing supported by a Medical Research Council (MRC) Clinical Infrastructure award (M008924) to J.M. The human embryonic and fetal material was provided by the MRC/Wellcome-Trust funded Human Developmental Biology Resource (HBDR, http://www.hdbr.org) (MR/R006237/1, MR/X008304/1, and 226202/Z/22/Z). Postnatal pancreas tissue was provided by the Bart’s Pancreas Tissue Bank (BPTB) and we acknowledge BPTP members Claude Chelala, Ahmet Imrali, Christine Hughes, and Tissue Access Chair Richard Grose. This study was also supported by the National Institute for Health and Care Research (NIHR) Exeter Biomedical research Centre (BRC). The view expressed are those of the author(s) and not necessarily those of the NIHR of the Department of Health and Social Care.

