## Supplementary Figures for "Developmentally dynamic changes in DNA methylation in the human pancreas"

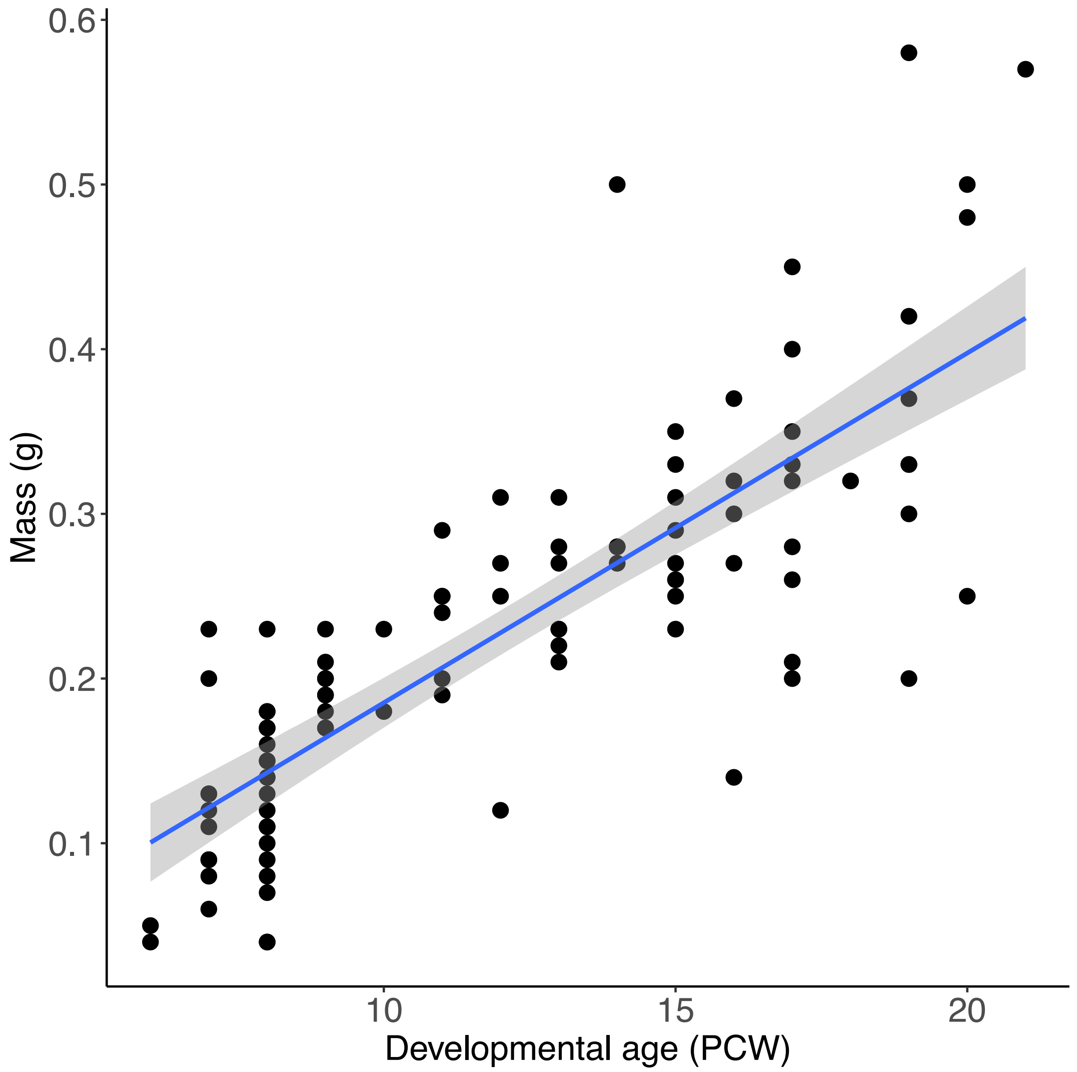


**Supplementary Figure 1. Fetal pancreas mass increases with developmental age.** Shown is a scatterplot of the mass (g) of each human fetal pancreas sample used in this study obtained from the HDBR showing a strong positive correlation with age (correlation = 0.803, $P < 2.2 \times 10^{-16}$).

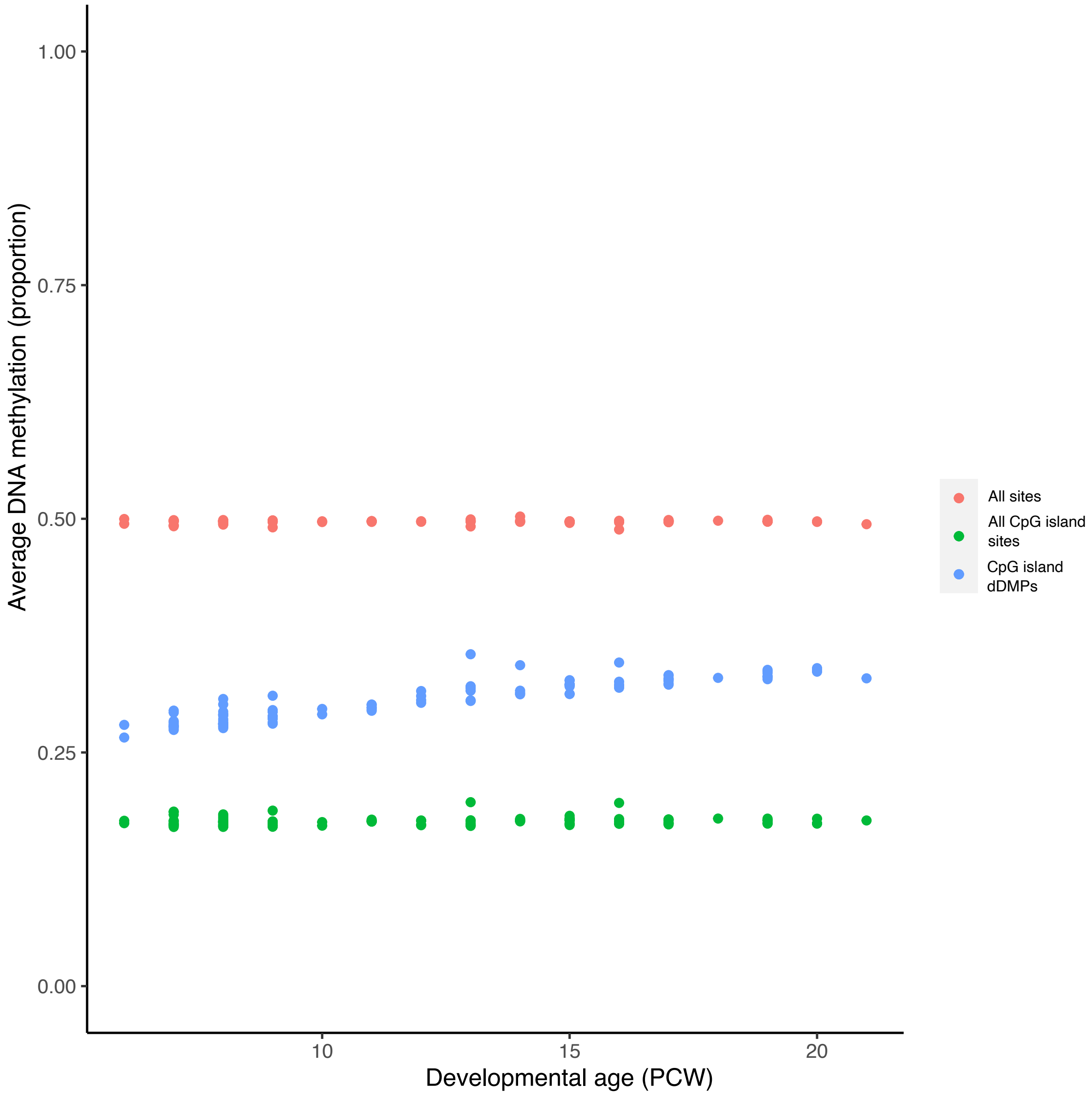


**Supplementary Figure 2. Global levels of DNA methylation remain stable across pancreas development.** There was no developmental change in mean DNA methylation across all sites tested (red, regression coefficient = 0.0055, $P = 0.497$) and also across all CpG island sites (green, regression coefficient = $8.62 \times 10^{-5}$, $P = 0.421$). In contrast CpG island dDMPs were characterized by an overall increase in DNA methylation across pancreas development (blue, regression coefficient = 0.0046, $P < 2.2 \times 10^{-16}$).

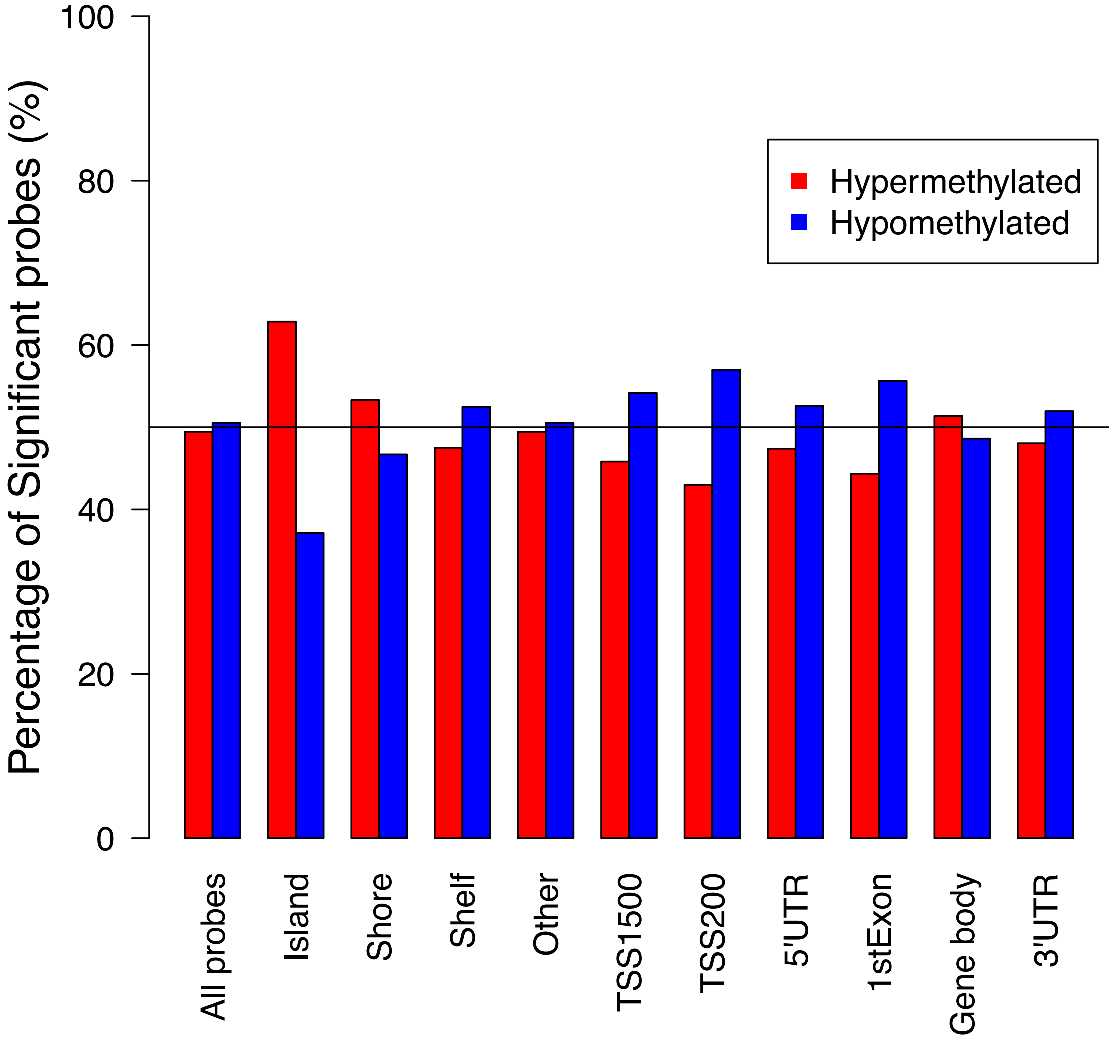


**Supplementary Figure 3. Proportion of hypermethylated and hypomethylated dDMPs across genic features.** Shown is the proportion of dDMPs that are characterized by either an increase in DNA methylation across development (red) or a decrease in DNA methylation (blue) split by genic region. There was an enrichment of hypermethylated dDMPs within CpG islands (62.9%), CpG shores (53.2%) and gene body (52.19%) regions, and an enrichment of hypomethylated dDMPs within TSS1500 (53.90%), TSS200 (59.88%) and 1$^{st}$ exon (58.25%) regions.

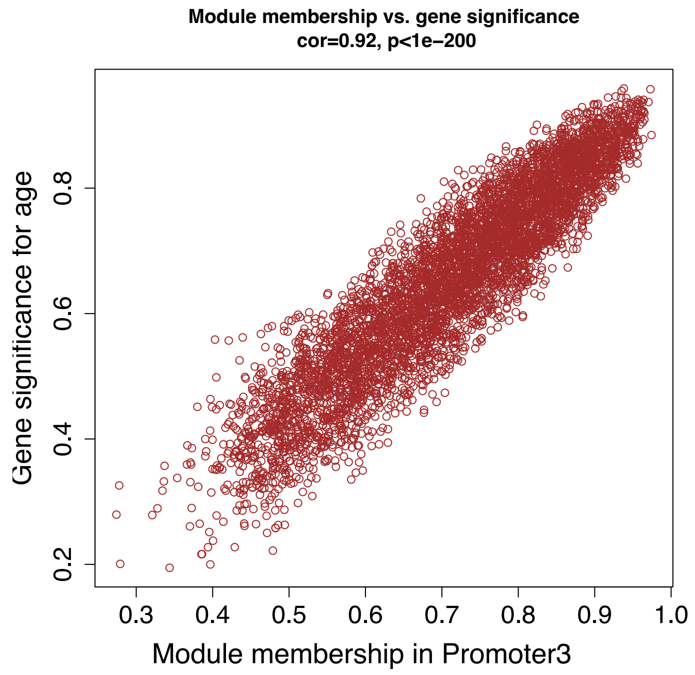


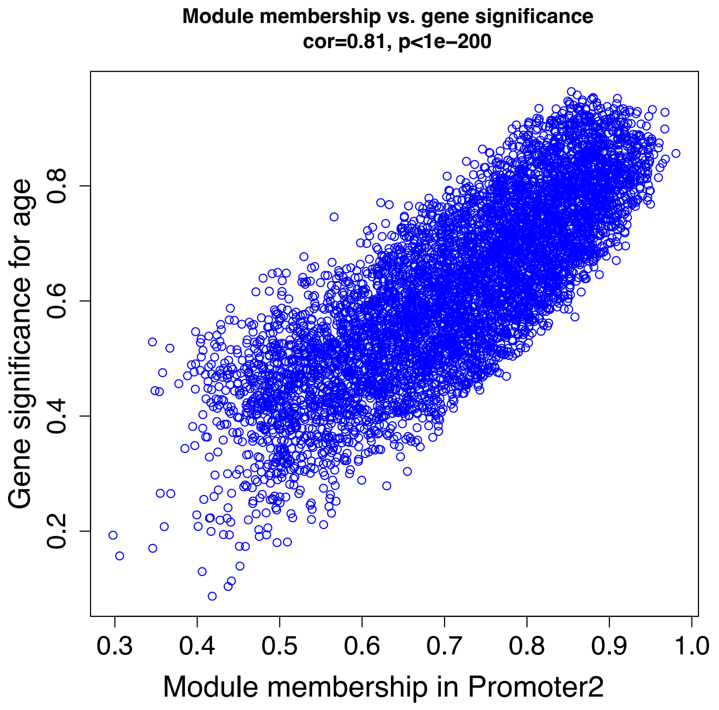


**Supplementary Figure 4. Module membership for DNA methylation sites in the Promoter3 (top) and Promoter2 (bottom) modules is highly correlated with site significance.**

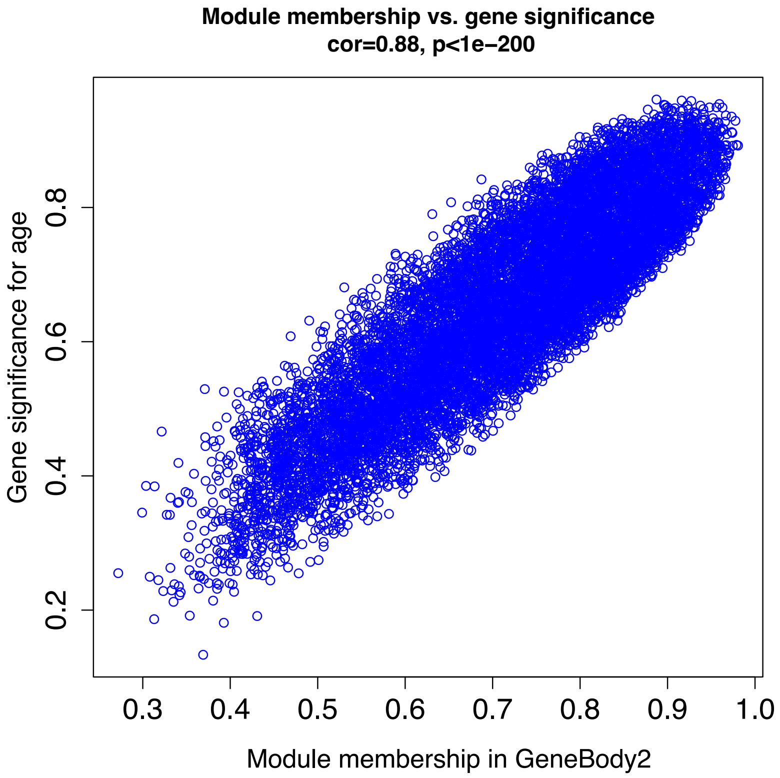


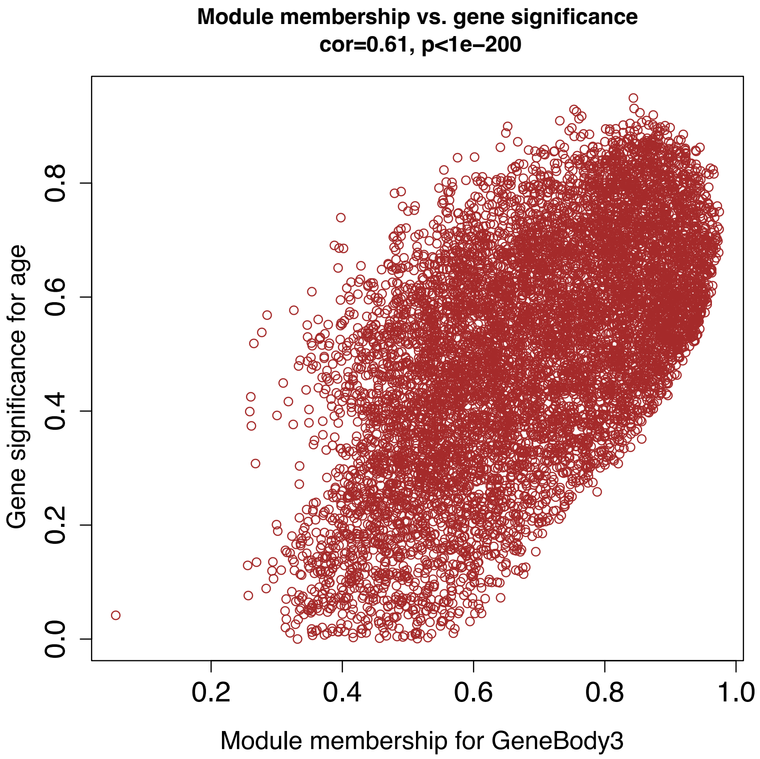


**Supplementary Figure 5. Module membership for DNA methylation sites in the GeneBody3 (top) and GeneBody4 (bottom) modules is highly correlated with site significance.**

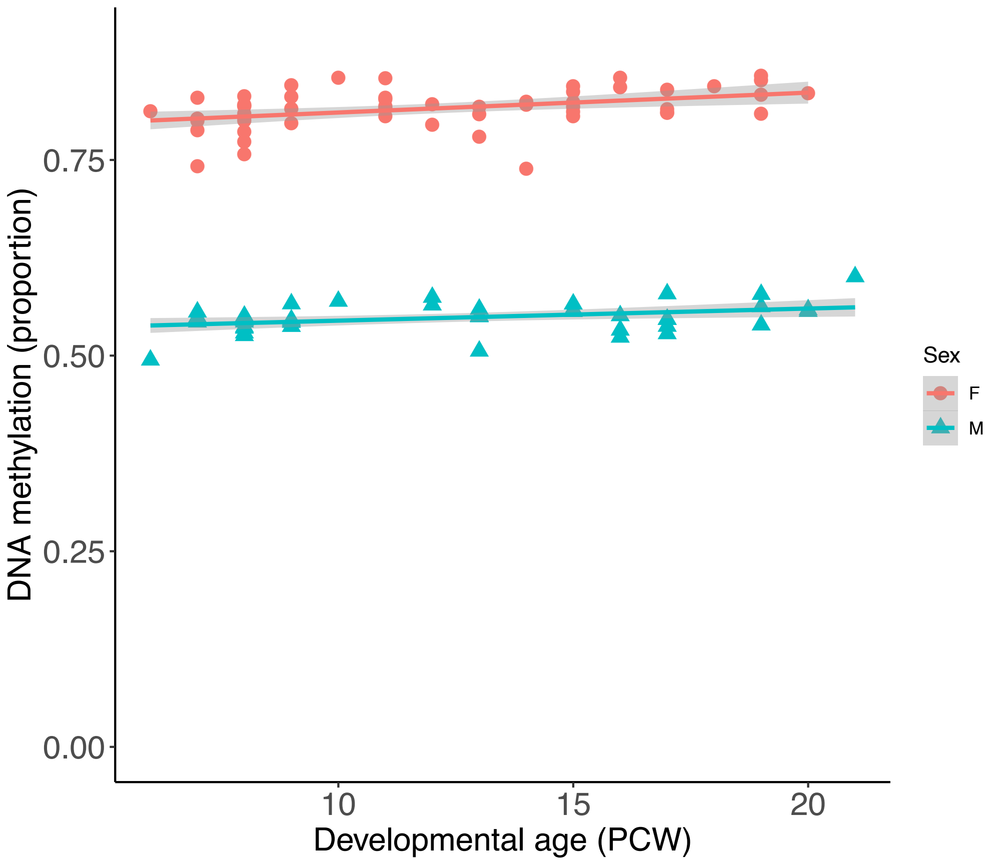


**Supplementary Figure 6. An example of an autosomal sex difference in DNA methylation in the developing pancreas.** The top-ranked autosomal DMP between males and females was cg06513015 ($P = 2.34 \times 10^{-78}$) which is annotated to the promoter region of the *ERV3-1* gene ($P = 2.34 \times 10^{-78}$). All autosomal sex differences in fetal pancreas DNA methylation are listed in **Supplementary Table 11**.

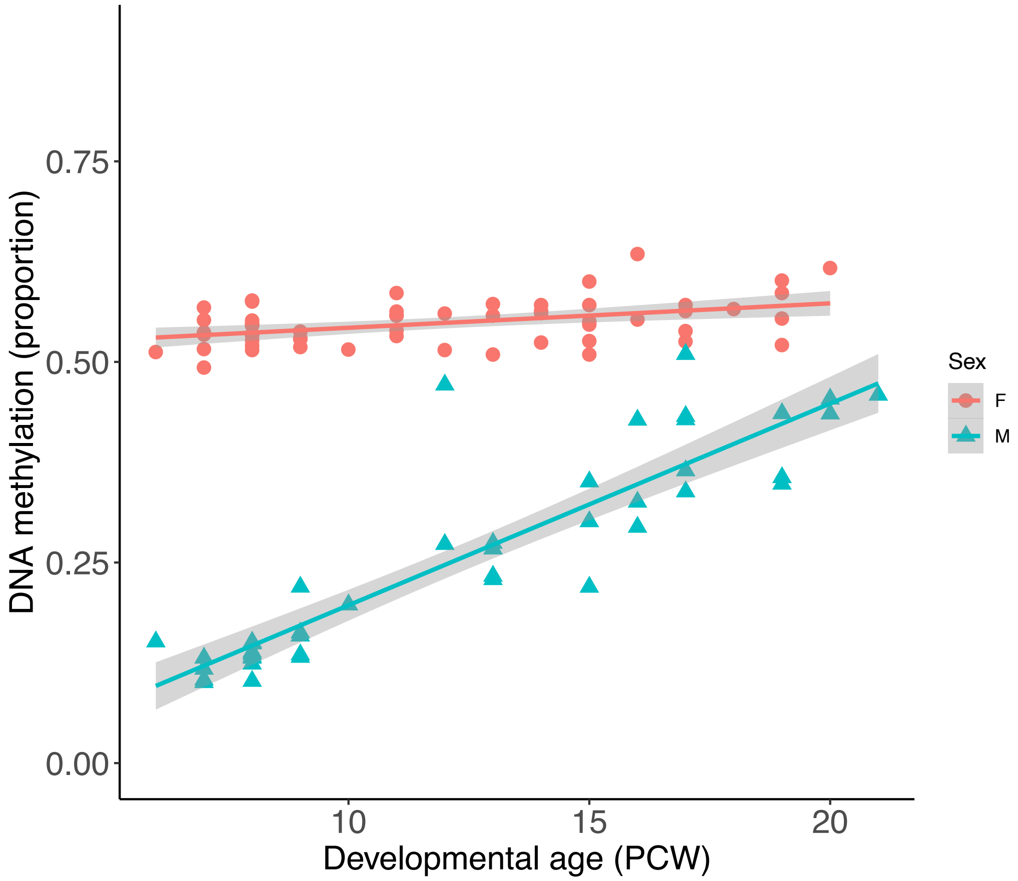


**Supplementary Figure 7. An example of a site on the X chromosome characterized by sex-specific changes in DNA methylation across pancreas development.** Shown is the top-ranked site (cg16354105) characterized by a sex-specific developmental trajectory in DNA methylation (P = 5.16E-20). All sites characterized by an interaction between sex and developmental age are listed in **Supplementary Table 12**.

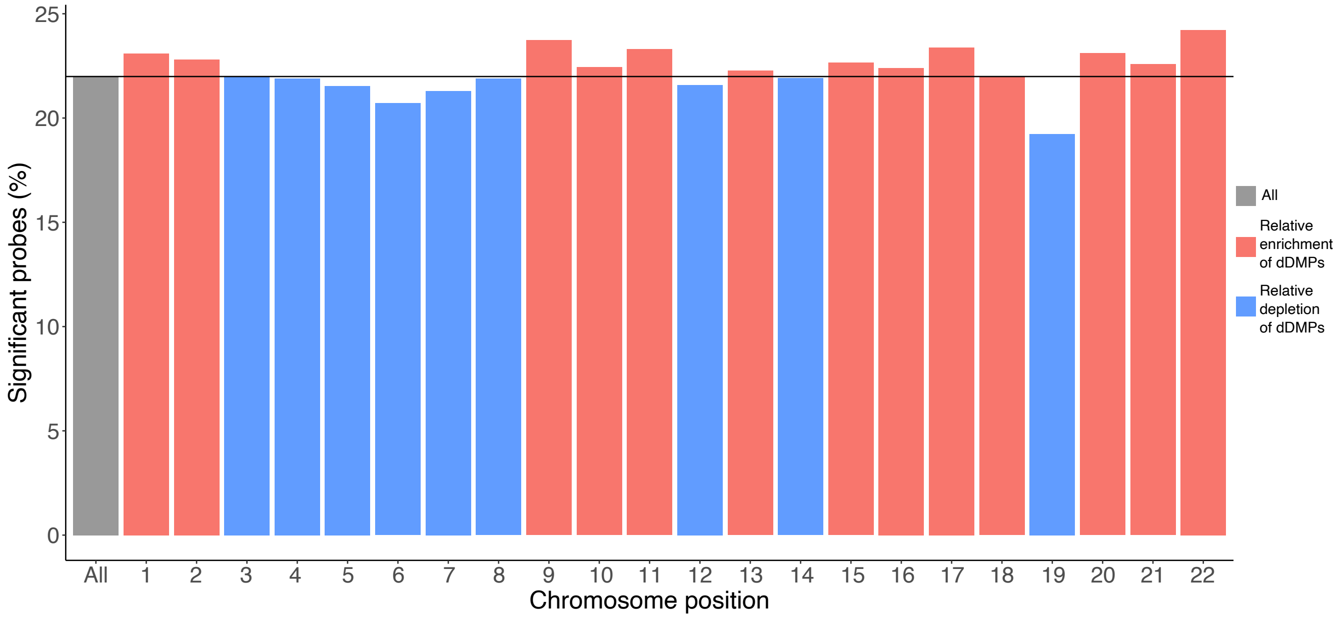


**Supplementary Figure 8. The distribution of fetal pancreas dDMPs is relatively consistent across autosomes**. Shown is the proportion of sites at which DNA methylation is significantly associated with pancreas development across all autosomes.

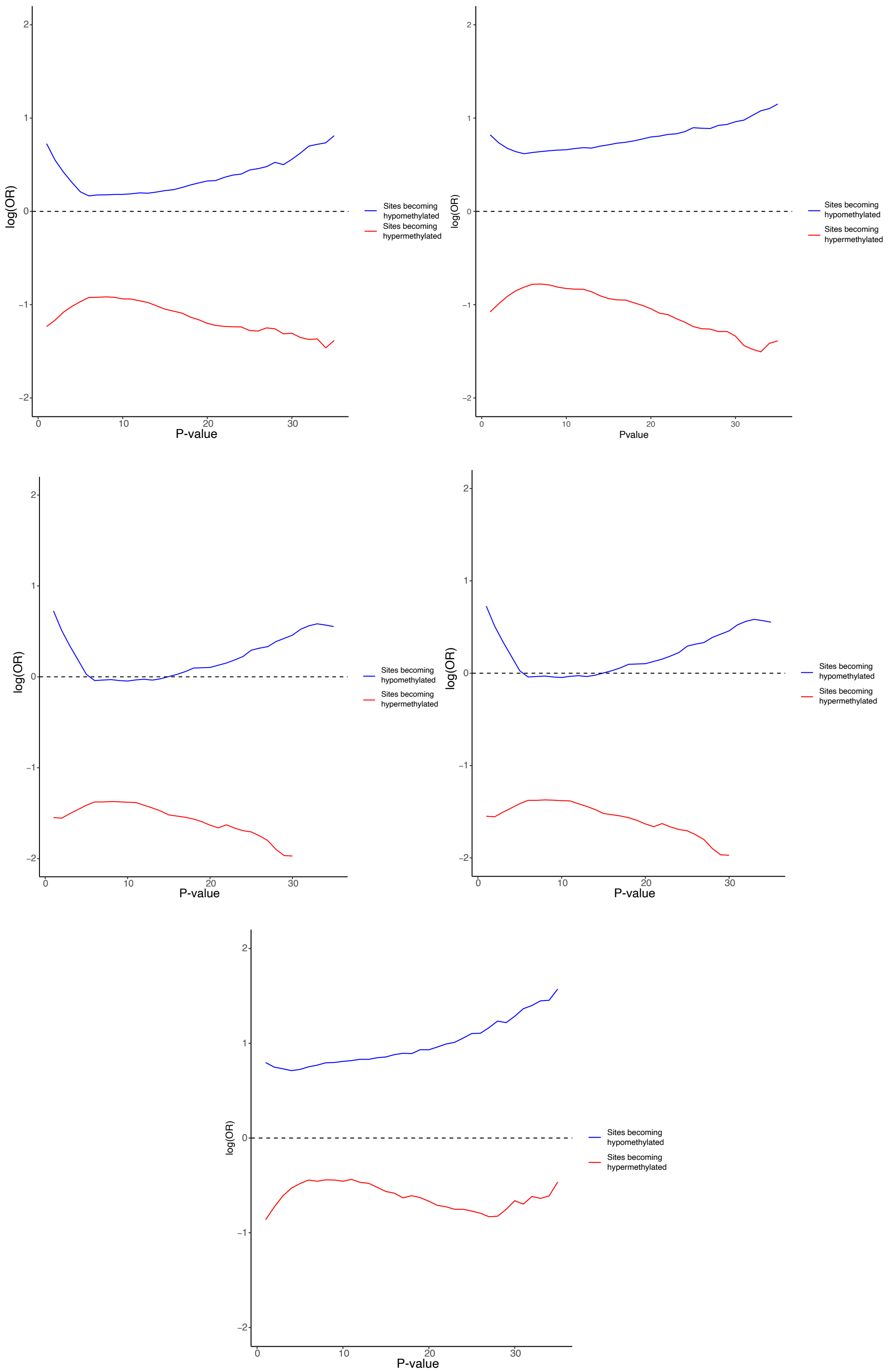


**Supplementary Figure 9**. **Relative enrichment of dDMPs in regions of transcription factor binding occupancy.** Shown is the enrichment of hypermethylated (red) and hypomethylated (blue) dDMPs in regulatory domains defined by key transcription factor binding site occupancy for key pancreatic transcription factors (PDX1, FOXA2, NKX6.1, NKX2.2 and MAFB).

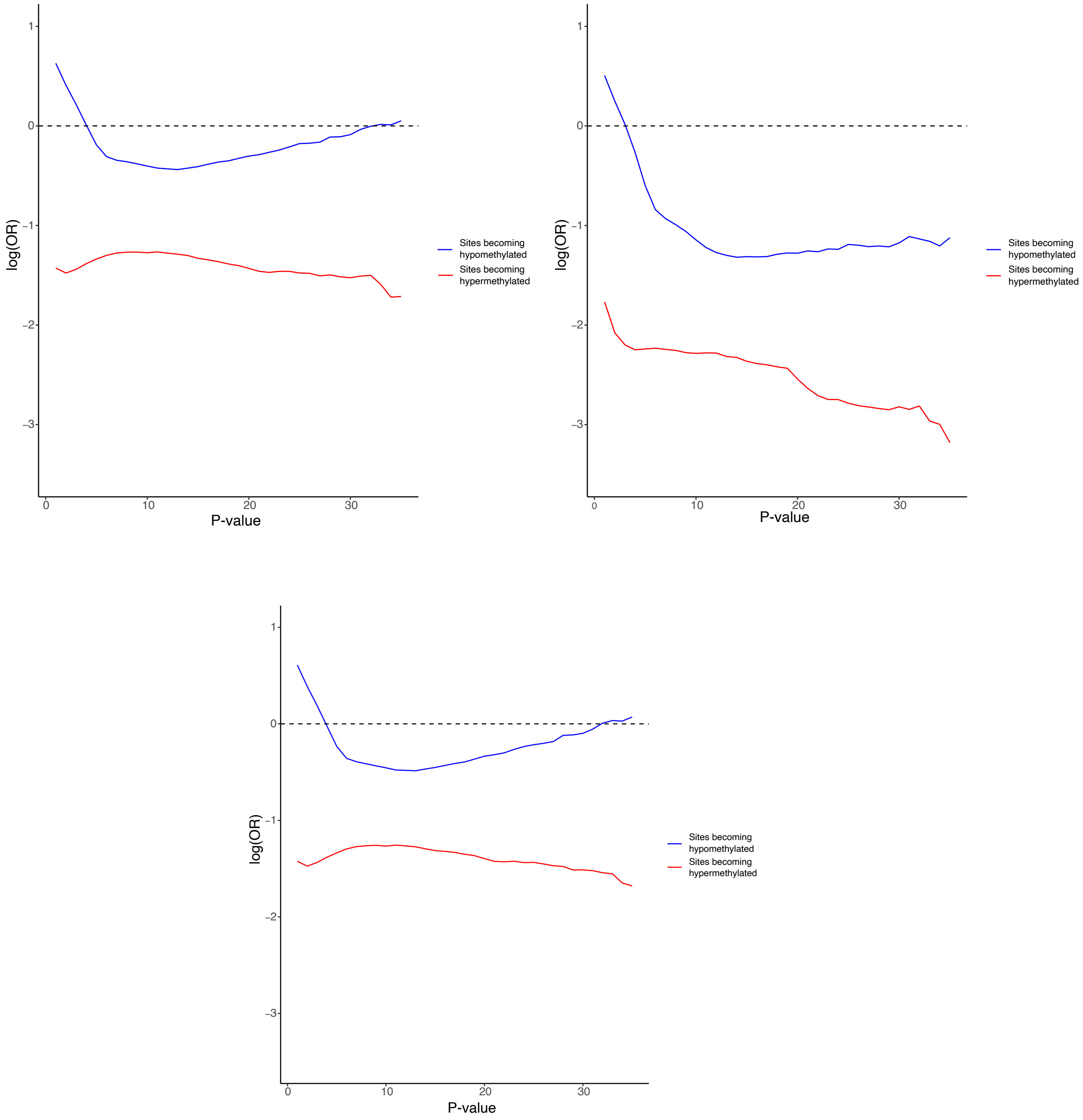


**Supplementary Figure 10**. **Relative enrichment of pancreas dDMPs in regions in regions of open chromatin.** Shown is the enrichment of hypermethylated (red) and hypomethylated (blue) dDMPs in genomic regions of defined by open chromatin in three endocrine cell types: (A) $\alpha$-cells, (B) $\beta$-cells, (C) $\delta$-cells.

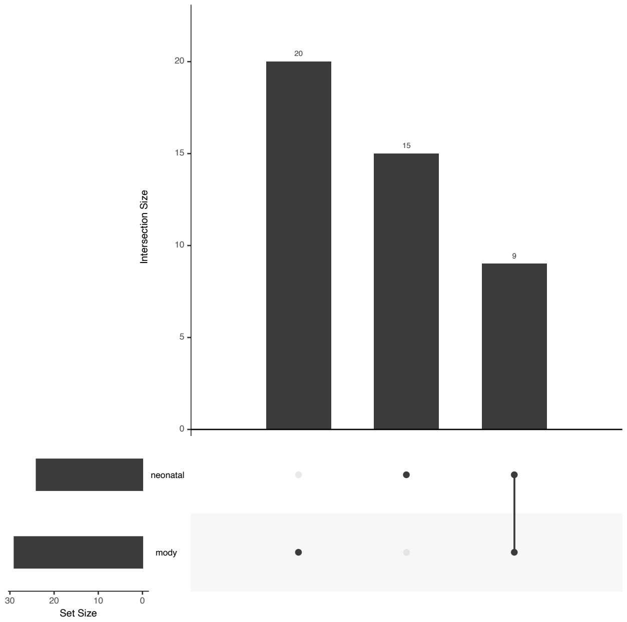


**Supplementary Figure 11. Overview of the neonatal diabetes and MODY genes tested for enrichment.** A full list of disease gene regions is given in **Supplementary Table 14** and **Supplementary Table 15**.

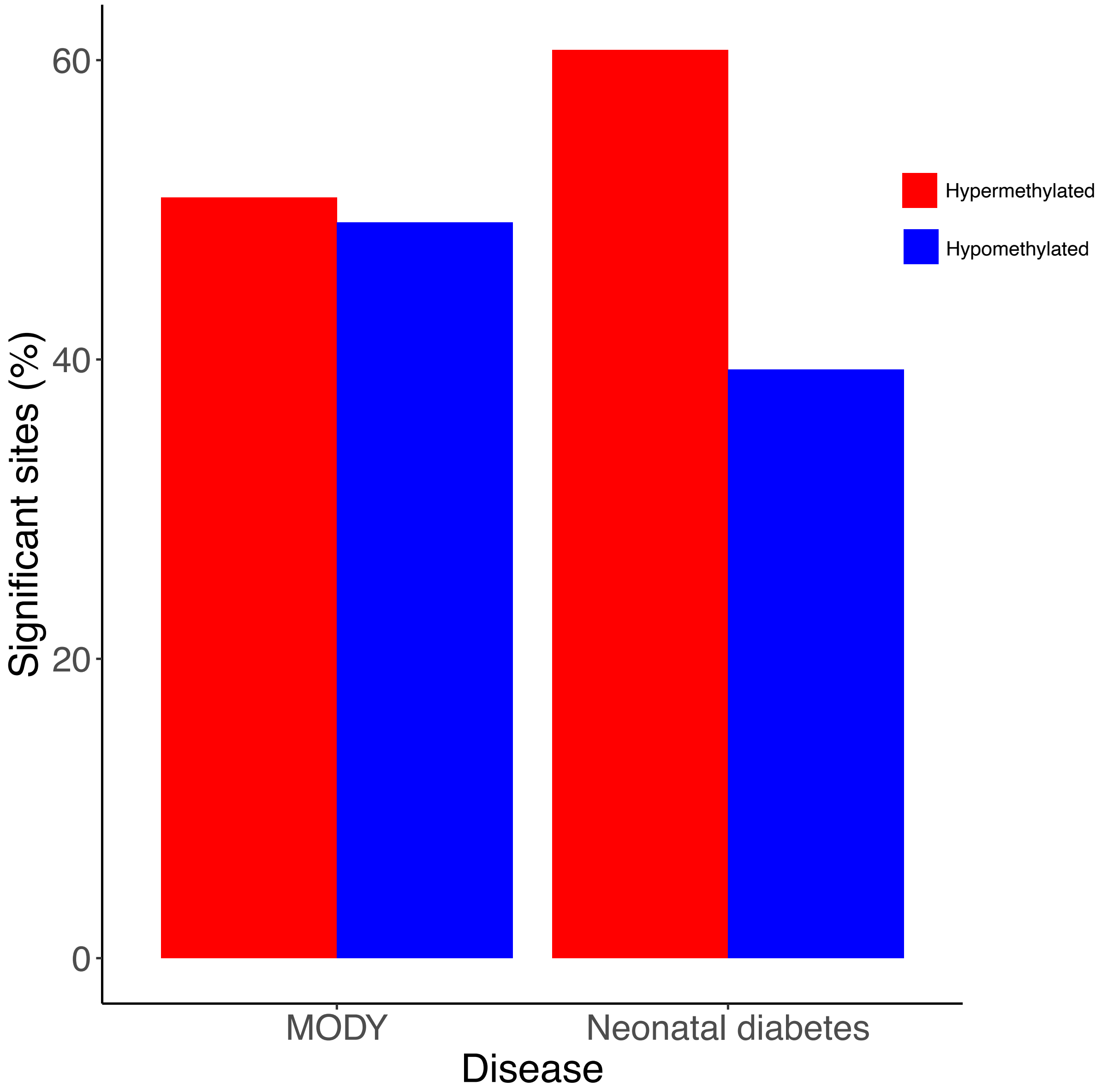


**Supplementary Figure 12. There is enrichment of dDMPs characterized by an increase in DNA methylation across development within genomic regions annotated to neonatal diabetes genes.**

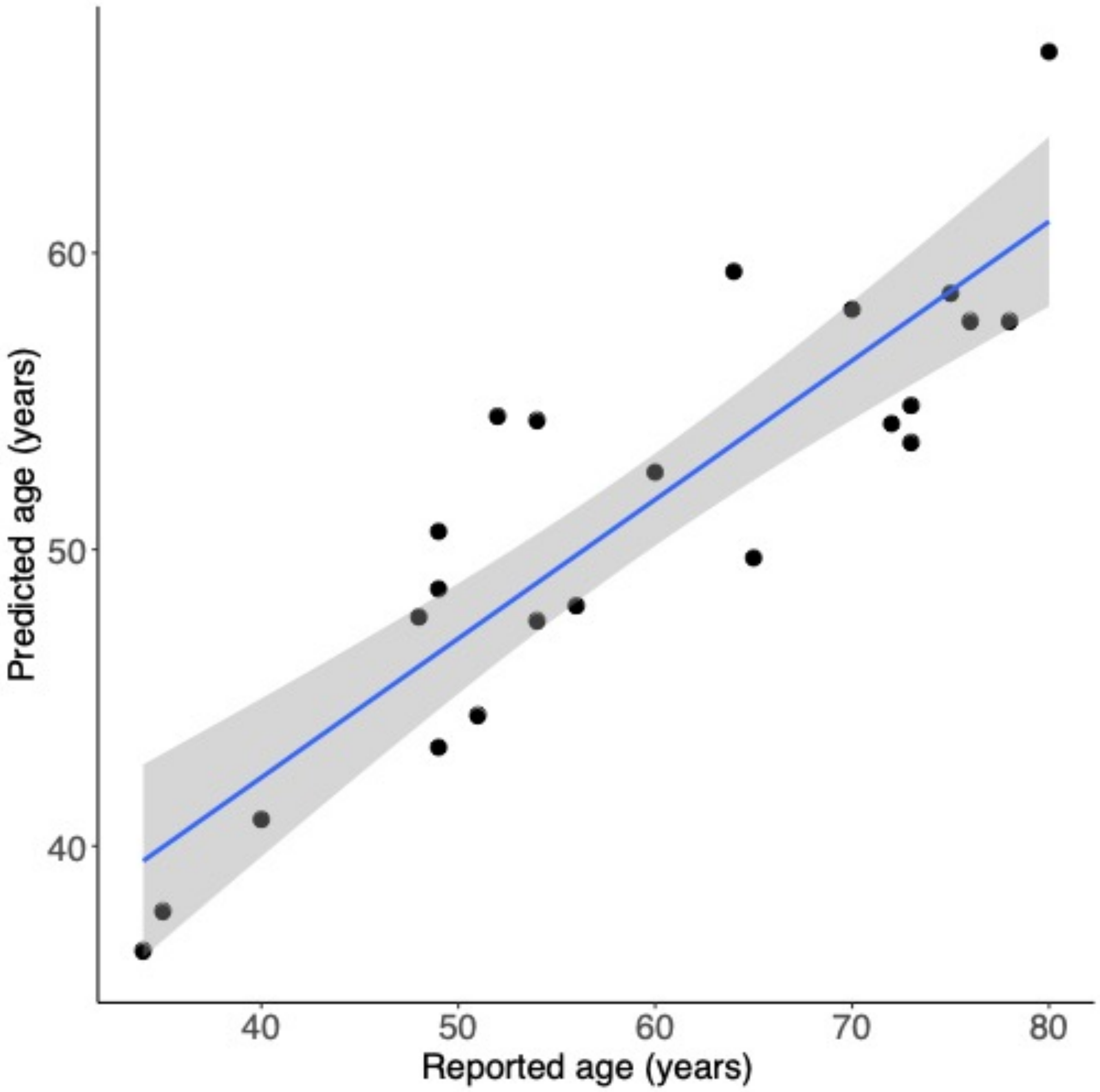


**Supplementary Figure 13. The chronological age of postnatal donors was strongly correlated with epigenetic age estimates derived from DNA methylation data.** Shown is the relationship between actual age and estimated age derived using the Horvath multi-tissue epigenetic clock (corr = 0.89, $P = 8.31 \times 10^{-35}$).

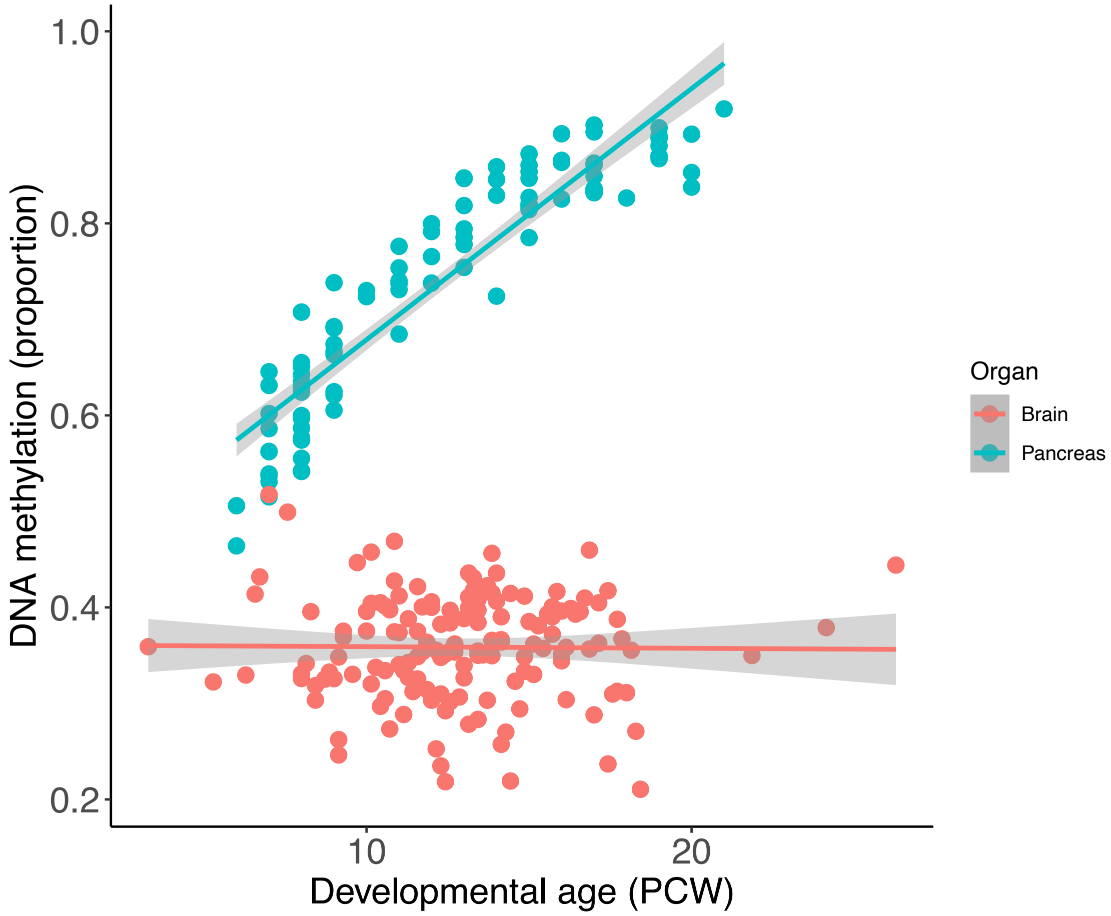


**Supplementary Figure 14. Pancreas specific developmental changes associated with *NNAT*.** Shown is the pancreas dDMP cg22551578 which displays dynamic change in DNA methylation in the developing pancreas (mean change in DNA methylation across developmental age = 0.025, $P$ = 5.18 x $10^{-36}$). In contrast, this site is characterized by no change in DNA methylation across development of the fetal brain (mean change in DNA methylation across developmental age = -0.00016, $P$ = 0.908).
