## Supplementary Table Legends for "Developmentally dynamic changes in DNA methylation in the human pancreas"

**Supplementary Table 1. Overview of all fetal and adult pancreas samples used in this study.** Shown for each sample is the tissue-bank ID, age, sex and Illumina EPIC array position.

**Supplementary Table 2. All significant dDMPs associated with human pancreas development.** Shown are all 177,130 dDMPs at which DNA methylation is significantly associated ($P < 9 \times 10^{-8}$) with pancreas development.

**Supplementary Table 3. GO analysis on genes annotated to the top 10,000 hypermethylated dDMPs.** Shown is the list of GO terms significantly enriched (FDR < 0.1) amongst genes annotated to hypermethylated dDMPs.

**Supplementary Table 4. KEGG analysis on genes annotated to the top 10,000 hypermethylated dDMPs.** Shown is the list of KEGG terms significantly enriched (FDR < 0.1) amongst genes annotated to hypermethylated dDMPs

**Supplementary Table 5. GO analysis on genes annotated to the top 10,000 hypomethylated dDMPs.** Shown is the list of GO terms significantly enriched (FDR < 0.1) amongst genes annotated to hypomethylated dDMPs.

**Supplementary Table 6. KEGG analysis on genes annotated to the top 10,000 hypomethylated dDMPs.** Shown is the list of KEGG terms significantly enriched (FDR < 0.1) amongst genes annotated to hypomethylated dDMPs.

**Supplementary Table 7. All significant dDMRs associated with human pancreas development.** Shown is the list of dDMRs which are significantly associated (corrected $P < 0.05$, number of sites ≥ 3) with pancreas development.

**Supplementary Table 8. Promoter region comethylation modules in the developing pancreas.** Shown are promoter WGCNA modules incorporating > 100 DNA methylation sites. All modules significantly associated with development are highlighted in grey.

**Supplementary Table 9. Gene body comethylation modules in the developing pancreas.** Shown are gene body WGCNA modules incorporating > 100 DNA methylation sites. All modules significantly associated with development are highlighted in grey.

**Supplementary Table 10. Pathway analysis of genes annotated to key members of comethylation modules associated with pancreas development.** Shown is all significant GO and KEGG pathways (FDR < 0.1) enriched amongst genes annotated to sites in significant promoter and gene body WGCNA modules.

**Supplementary Table 11. Autosomal DMPs associated with sex in the human fetal pancreas.** Shown is a list of autosomal sites at which DNA methylation is significantly different ($P < 9 \times 10^{-8}$) between males and females in the human fetal pancreas.

**Supplementary Table 12. DNA methylation sites characterized by sex specific changes in the human fetal pancreas**. The list of dDMPs with significant ($P = 9 \times 10^{-8}$) sex-specific changes across pancreatic development.

**Supplementary Table 13. Enrichment statistics for all genomic and regulatory regions.** Shown are the enrichment statistics at genome-wide significance threshold ($P < 9 \times 10^{-8}$).

**Supplementary Table 14. Overview of dDMPs associated with MODY genes.**

**Supplementary Table 15. Overview of dDMPs associated with neonatal diabetes genes.**

**Supplementary Table 16. All dDMPs associated with neonatal diabetes genes.**

**Supplementary Table 17. A comparison of developmental changes in pancreas and brain DNA methylation at pancreas dDMPs.** All significant pancreas dDMPs ($9 \times 10^{-8}$) also present in the fetal brain.

**Supplementary Table 18. A comparison of developmental changes in pancreas and brain DNA methylation at brain dDMPs.** All significant brain dDMPs ($9 \times 10^{-8}$) also present in the fetal pancreas.

**Supplementary Table 19. All pancreas dDMPs with no corresponding developmental changes in the brain.**

**Supplementary Table 20. All brain dDMPs with no corresponding developmental changes in the pancreas.**

**Supplementary Table 21. Reagents used for pancreatic staining.** A) Details of the primary antibodies used for immunohistochemical staining. B) Details of antibodies and conditions required for immunofluorescence staining. C) Details of all other accessory agents used for both staining protocols.
